# AlphaFold as a Prior: Experimental Structure Determination Conditioned on a Pretrained Neural Network

**DOI:** 10.1101/2025.02.18.638828

**Authors:** Alisia Fadini, Minhuan Li, Airlie J. McCoy, Thomas C. Terwilliger, Randy J. Read, Doeke Hekstra, Mohammed AlQuraishi

## Abstract

Advances in machine learning have transformed structural biology, enabling swift and accurate prediction of protein structure from sequence. However, challenges persist in capturing sidechain packing, condition-dependent conformational dynamics, and biomolecular interactions, primarily due to scarcity of high-quality training data. Emerging techniques, including cryo-electron tomography (cryo-ET) and high-throughput crystallography, promise vast new sources of structural data, but translating experimental observations into mechanistically interpretable atomic models remains a key bottleneck. Here, we address these challenges by improving the efficiency of structural analysis through combining experimental measurements with a landmark protein structure prediction method – AlphaFold2. We present an augmentation of AlphaFold2, ROCKET, that refines its predictions using cryo-EM, cryo-ET, and X-ray crystallography data, and demonstrate that this approach captures biologically important structural variation that AlphaFold2 does not. By performing structure optimization in the space of coevolutionary embeddings, rather than Cartesian coordinates, ROCKET automates difficult modeling tasks, such as flips of functional loops and domain rearrangements, that are beyond the scope of current state-of-the-art methods and, in some instances, even manual human modeling. The ability to efficiently sample these barrier-crossing rearrangements unlocks a new horizon for scalable and automated model building. Crucially, ROCKET does not require retraining of AlphaFold2 and is readily adaptable to multimers, ligand-cofolding, and other data modalities. Conversely, our differentiable crystallographic and cryo-EM target functions are capable of augmenting other structure prediction methods. ROCKET thus provides an extensible framework for the integration of experimental observables with biomolecular machine learning.

## 1 Introduction

Machine learning (ML) has revolutionized structural biology by enabling highly accurate protein structure prediction. Breakthrough models such as AlphaFold2 (AF2) [1], RoseTTAFold [2], and their descendants harness coevolutionary signals in large-scale sequence data to produce predictions with atomic-level precision and near-experimental accuracy [3, 4]. Despite this accomplishment, computational approaches still struggle to capture important properties such as sidechain packing, functional dynamics, and large-scale molecular assembly [5–9].

The success of these ML models relies heavily on a vast collection of experimentally resolved structures – made possible by decades of effort by the structural biology community. However, high-quality ground-truth data that capture multiple functional states, biomolecular interactions, and structural variations are scarce. Advances in high-throughput experimental techniques are beginning to provide such data. For example, modern crystallography beamlines and single-particle cryo-electron microscopy (cryo-EM) can now yield datasets under many perturbations, e.g. for drug screening, and tracking of structural changes during biochemical transformations [10, 11], while cryo-electron tomography (cryo-ET) enables *in situ* observations of macromolecular complexes [12, 13]. These advances promise deeper insights into conformational flexibility and complex assembly.

A key bottleneck in processing these large datasets is the reconstruction of atomic models from experimental observations. ML methods have helped streamline this process by providing high-quality starting points for atomic model building [6, 14, 15]. Standard refinement software [16–18] improves these starting points by optimizing experimental likelihoods in Cartesian coordinate space, complemented by pattern matching-based model building to overcome local barriers [19–21]. This combination struggles when the structural rearrangements presented by the data are large, such as switches in secondary structure, flips in flexible loops, or shifts in relative domain orientations. Model building becomes particularly challenging, even for humans, below 4–5 Å resolution, where critical structural details such as sidechains become indistinct. This difficulty is especially pronounced in the emerging field of cryo-ET, which currently tends to produce data at very low resolution [22]. New priors that guide model building could therefore yield more accurate atomic models for low-resolution datasets and limit labor-intensive manual intervention when initial structures deviate from the experimental data.

We hypothesized that the implicit prior structural knowledge embedded in pre-trained ML structure prediction methods could guide atomic model building from experimental data more efficiently than traditional geometric restraints [23]. Historically, the integration of model and experiment was facilitated by the explicitly physics-based nature of structure prediction methods, which readily permitted the incorporation of experiment-derived potential functions [24, 25]. “Black-box” ML models like AF2, however, make this integration less straightforward. Recent efforts to adapt ML-based structure prediction to incorporate experimental constraints have done so either through fine-tuning of weights from existing architectures [26, 27] or full retraining, as exemplified by the ModelAngelo method for cryo-EM data [28]. These strategies are promising but computationally expensive, data-hungry, and limited to the data modality they were trained on. Another approach, PredictAndBuild [6, 29], iterates between predicting structure conditioned on a template, and rebuilding the predicted structures based on experimental data to yield the next template. This approach avoids modifying the structure prediction method, but its decoupled prediction and rebuilding steps can work against each other and hinder convergence.

Ideally, existing ML methods could serve as implicit structural priors – without retraining – to accelerate and automate atomic model building. Indeed, the ColabDock framework incorporates cross-linking experimental restraints in protein complex structure prediction [30], and other contemporaneous works have begun exploring this direction for cryo-EM and crystallography, but their applicability to challenging modeling tasks remains unclear [31, 32]. Here, we combine the high accuracy of pre-trained AF2 structure prediction with guidance from experimental data in a way that can be flexibly adapted to different data modalities. We achieve this through ROCKET (Refining OpenFold with Crystallographic/Cryo-EM liKElihood Targets), a framework that integrates Open-Fold [33], a trainable re-implementation of AF2, with differentiable crystallographic and cryo-EM likelihood targets [34]. ROCKET refines structures at inference time (no retraining), steering predictions toward experimentally supported conformations (Fig. 1a). Inspired by strategies for expanding AF2 to conformational sampling [35–39], we directly optimize the embedded multiple sequence alignment (MSA) cluster profile [36, 37] in AF2, transforming structure refinement to a data-guided search within AF2’s continuous representation of sequences.

**Fig. 1.**
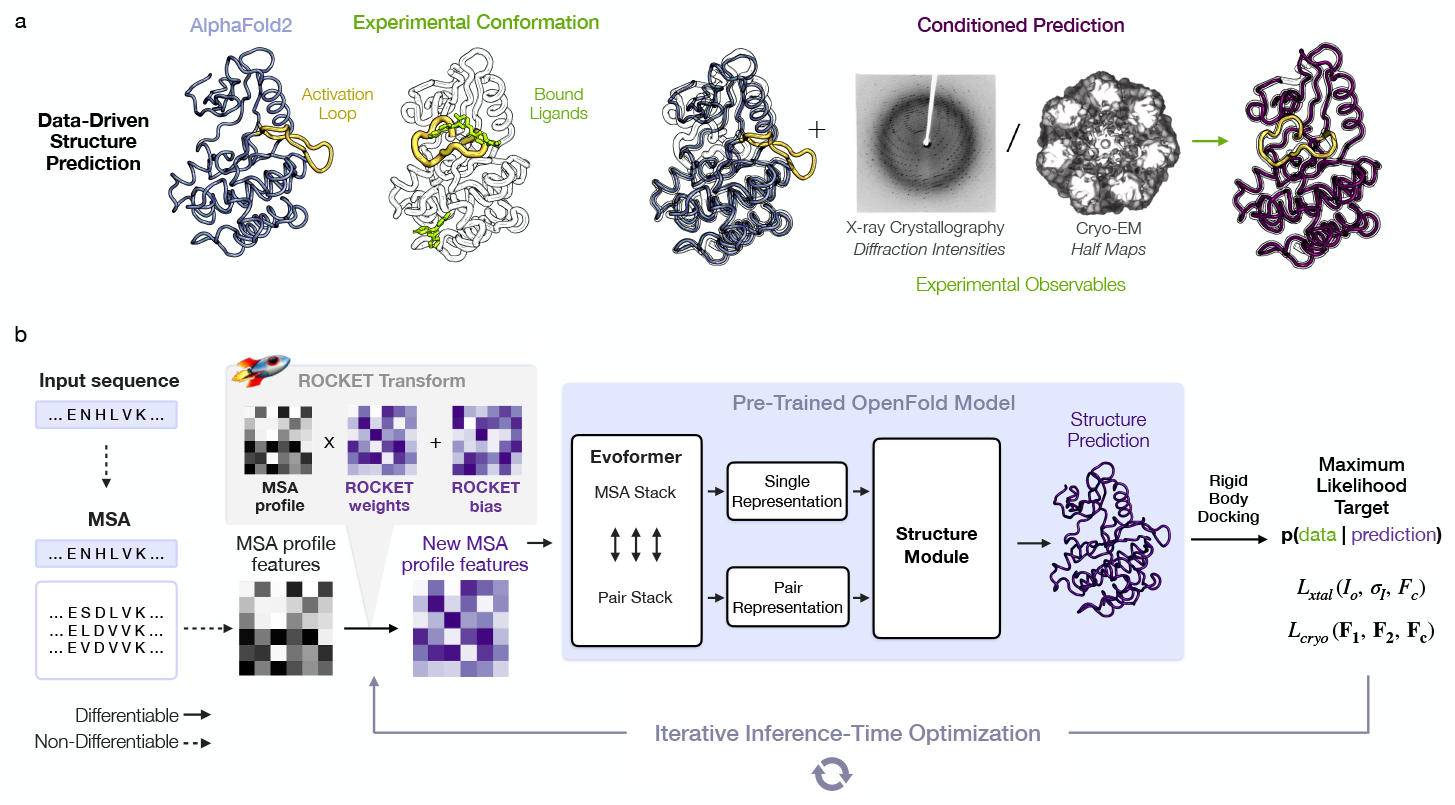
Data-Driven Structure Prediction Refinement with ROCKET. (a) Experimental techniques such as X-ray crystallography and cryo-EM provide high-fidelity observations of conformational states, capturing, for example, ligand-induced changes (right; activation loop of human c-Abl kinase trapped in the active state by a small-molecule activator – PDB ID 3PYY) that may not be modeled by ML-based predictions (left; AF2 prediction of c-Abl in the untrapped state). (b) ROCKET extends OpenFold by integrating crystallographic and cryo-EM likelihood targets within its differentiable prediction pipeline. It accomplishes this by learning, at inference-time, multiplicative and additive adjustments to MSA cluster profiles that maximize their agreement with experimental data as computed by an experimental likelihood function. For the crystallographic target, the function (*L*_xtal_) depends on observed crystallographic diffraction intensities and their measurement errors (*I*_*o*_, *σ*_*I*_) and on the structure factor amplitudes computed from the predicted model (*F*_*c*_). For the cryo-EM target, the function (*L*_cryo_) depends on the complex Fourier terms from experimental half-maps (**F**_**1**_ and **F**_**2**_) and the complex structure factors computed from the predicted model (**F**_**c**_).

Cryo-EM and crystallography are the predominant techniques for producing atomic-level insights into conformational heterogeneity, macromolecular interactions, and functional structural rearrangements. Despite crystallography’s promise for high-resolution, high-throughput screening of experimental conditions [40–42] and drug candidates [43], ML applications in protein crystallography [34, 44] lag behind cryo-EM [28, 45–47] due to challenges presented by crystal symmetry and phase loss. ROCKET integrates both X-ray and cryo-EM data directly into OpenFold’s inference process. We find that performing optimization in the latent MSA profile embedding, instead of in Cartesian coordinate space, allows for barrier-crossing structural rearrangements – such as domain shifts and drug-induced conformational changes – that conventional refinement methods struggle to access. Our evaluation of ROCKET on crystallographic test cases demonstrates that it can reliably capture backbone and sidechain conformations with accuracy comparable to that of manually curated models. Moreover, subsampling MSAs to generate alternative MSA profile embeddings improves initial predictions, helping to overcome limitations of gradient-descent in challenging cases.

ROCKET is particularly valuable for the unsolved challenge of model building at very low resolution [48], which is important for cryo-ET. Existing software struggles at resolutions worse than 4–5 Å: ModelAngelo was not trained to work with maps past 4 Å [28] and PredictAndBuild will not automatically rebuild with maps worse than 3.5 Å [6]. We find that ROCKET’s inference-time refinement allows it to explore a wide conformational space and that it remains robust for atomic modeling in noisy, low-resolution maps, sometimes outperforming expert manual refinement. Since ROCKET relies on an input sequence for AF2 inference, it is complementary to other ML approaches designed to identify proteins in cryo-EM maps [28, 49, 50]. Its capabilities render ROCKET a generalizable approach for integrating experimental data with ML-based biomolecular modeling.

## 2 Results

### 2.1 Method Overview

AF2-based structure prediction begins with the construction of a multiple sequence alignment (MSA), typically through a search of sequence databases for proteins homologous to the query of interest. The resulting MSA comprises aligned sequences of identified homologs, from which one can infer a protein’s evolutionary history, including its patterns of residue-residue coevolution. AF2 transforms raw MSAs into an input representation suitable for neural network computation, known as the MSA cluster profile. Through many studies and ablations, it has been shown that the depth and diversity of the MSA, and the statistical patterns found therein, determine the geometry and quality of a structure predicted by AF2 [1]. Given this central role, we reasoned that direct optimization of the continuous space of MSA cluster profiles would provide the greatest lever for influencing AF2 predictions; a hypothesis supported by recent observations made by other groups [36, 37] as well as our own experiments (Fig. S1).

To operationalize this principle, ROCKET augments OpenFold with a new module that optimizes MSA cluster profiles to maximize agreement between predictions and experimental observables (Fig. 1b; see Methods for algorithmic details). Through an experimental target function that quantifies this agreement, ROCKET performs gradient descent in the space of MSA cluster profiles. During each descent step, ROCKET computes a forward pass through OpenFold, evaluates the target function and its derivatives, then updates the MSA cluster profile to increase the function’s agreement with model predictions. ROCKET currently provides two target functions, for crystallographic and cryo-EM data, and is extensible to other data modalities.

Since OpenFold-predicted structures are generated in an arbitrary reference frame, ROCKET first performs molecular replacement [51] or cryo-EM docking [52] to align the predicted model with the experimental data before starting the iterative refinement process. The resulting roto-translation is applied at every subsequent iteration to align the model. While only approximately optimal, previous observations indicate that initial AF2 predictions are usually sufficiently accurate for robust placement in the data [14, 15, 29, 53]. After iterative refinement through OpenFold is complete, a final local structure refinement is performed using phenix.refine [16] to optimize local geometry and atomic displacement parameters. Owing to memory limitations, ROCKET currently only operates on one protein chain at a time and requires special handling for cases involving multiple chains (see “Multi-Chain Dataset Handling” in Methods).

### 2.2 Evaluation Dataset and Approach

We took a two-pronged approach to assessing ROCKET’s effectiveness in guiding structure prediction with experimental data. In the first, we focused on minor refinements and high-resolution datasets that are already well-served by existing, conventional methods, and where ROCKET’s primary utility would be in streamlining experimental model building through integration with ML-based structure prediction. Here, our focus was on matching the best methods on relatively easy problems. In the second and more ambitious prong, we tackled cases where existing automated methods break down, either due to the magnitude and complexity of the required rearrangements or the low resolution of the data; cases in which manual human intervention is necessary and may not even be sufficient.

For the first prong, we selected a diverse set of 27 high-resolution, single-chain X-ray crystallographic datasets and their corresponding deposited structures (Fig. S2). We prioritized crystallography because, as noted earlier, its integration with ML-based methods remains underdeveloped [34]. All 27 datasets were solved at resolutions better than 3 Å (Table 1), a regime where conventional methods, such as phenix.refine, perform well and hybrid methods, such as PredictAndBuild, perform extremely well [29].

For the second prong, we examined large-scale structural changes in ligand-induced loop rearrangements in three human proteins: c-Abl kinase (PDB ID 3PYY), protein tyrosine phosphatase 1B (PTP-1B, PDB ID 1NWL), and the serpin plasminogen activator inhibitor-1 (PAI-1, PDB ID 1JL5). We also examined a cryo-EM structure of *E. coli* GroEL (PDB ID 9C0B) where the subunits adopt a markedly different conformation from the original AF2 prediction. Finally, to assess performance in the low-resolution regime, we selected a 3.82 Å crystallographic dataset of the human multidomain protease inhibitor (HAI-1, PDB ID 5H7V) and a 9.60 Å sub-tomogram average of *E. coli* GroEL (PDB ID 8P4P).

### 2.3 ROCKET Reliably Refines High-Resolution Structures

At high resolution, we expect deposited backbone and sidechain coordinates to serve as reliable ground truths for evaluation. To validate ROCKET’s performance on the 27 crystallographic structures in our first prong, we compared its refined models to those from the PDB-REDO database [54], which systematically re-refines and validates human-deposited coordinates. Fig. 2a shows C*α* root-mean-squared deviations (RMSDs) with respect to PDB-REDO models for original AF2 predictions and ROCKET-refined ones. ROCKET improves all AF2 predictions except one, bringing them closer to the experimentally determined structures. Focusing on the ten most difficult cases, where AF2 predictions deviate by more than 1 Å from the PDB-REDO models, ROCKET achieves substantial structural corrections (Fig. 2a, stars; mean RMSD drop of 0.47 Å), demonstrating robustness in challenging scenarios.

**Fig. 2.**
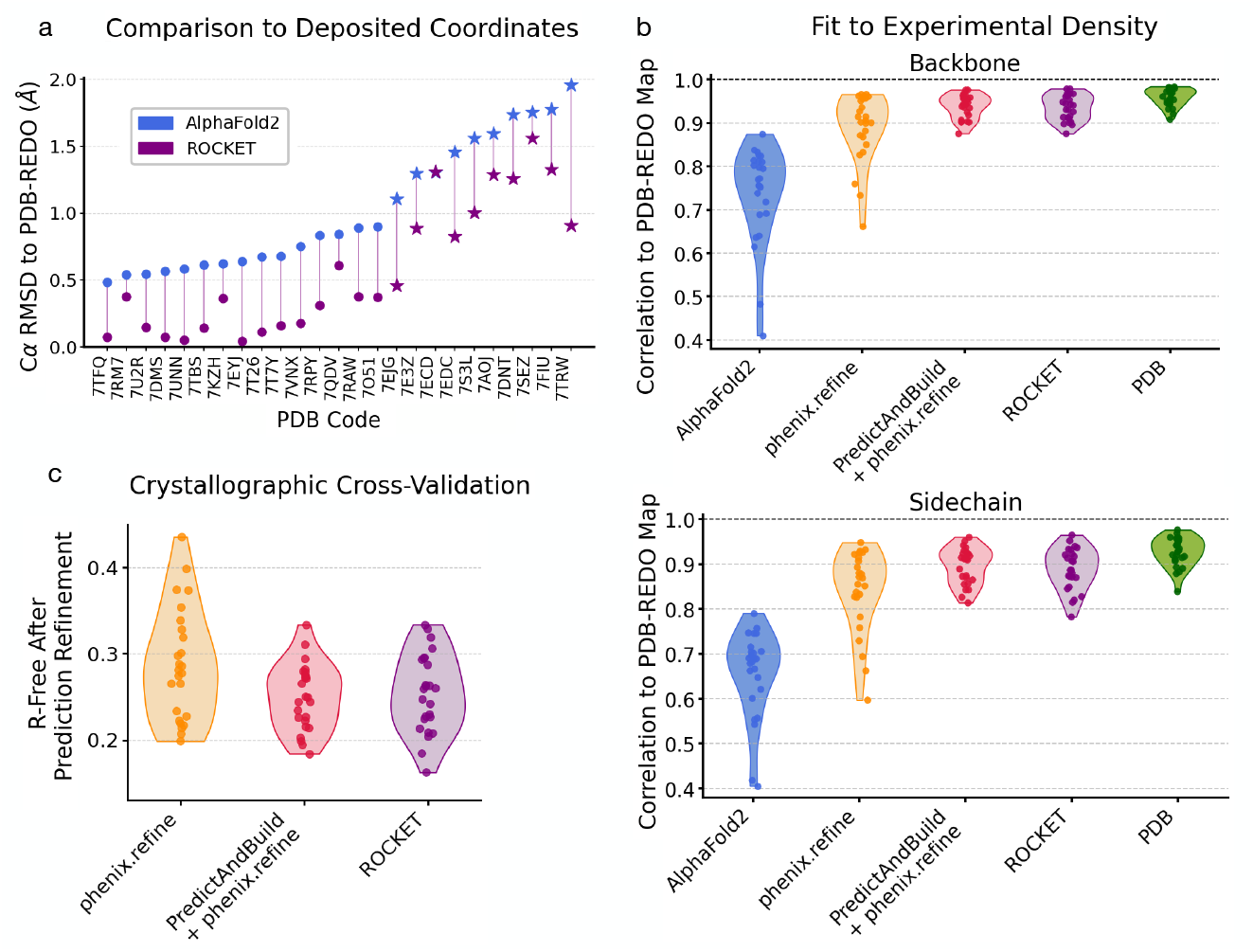
ROCKET Refines a Diverse Set of Predictions Using High-Resolution X-ray Crystallography Datasets. (a) C*α* RMSDs of conventional AF2 and ROCKET-refined predictions with respect to “ground truth” models from the PDB-REDO database, which systematically re-refines and validates human-deposited coordinates. AF2 predictions deviating by more than 1 Å from PDB-REDO models are highlighted with a star and generally correspond to instances where ROCKET performs the largest structural corrections. (b) Real-space Pearson correlation coefficient (RSCC) values of electron density maps derived from model structures vs. experimental amplitudes and unbiased phases from PDB-REDO. PDB-REDO maps are expected to have a favorable phase bias for deposited PDB models, artificially increasing their reported RSCCs. (c) Crystallographic R-free values for models placed by molecular replacement and refined using different methods.

Although C*α* RMSD is a convenient measure of overall model quality, it is prone to underestimating discrepancies between atomic models and data, especially in flexible loops or sidechains that exhibit low occupancy or disorder. For a more sensitive assessment, we employed the real-space Pearson correlation coefficient (RSCC), which directly compares model-derived electron density maps to maps from experimental amplitudes (here, combined with phases from PDB-REDO models). As shown in Fig. 2b, ROCKET substantially improves RSCC values for both backbones and sidechains across all test cases. Our automated rebuilding workflow achieves accuracies comparable to human-deposited models and performs on par with PredictAndBuild. Notably, optimizing MSA cluster profiles not only corrects secondary structures but also improves sidechain fit to the data (Fig. S4). Further validation using crystallographic R-free factors [55] (Fig. 2c) confirms ROCKET’s added value: all but two models see improvements relative to phenix.refine alone, with 10 models showing R-free reduction of more than 3%. ROCKET’s performance on R-free is comparable to the more complex combination of PredictAndBuild + phenix.refine, and the two can complement one another (Fig. S5).

We also performed an important control experiment: AF2 includes built-in confidence metrics, which several studies have leveraged to explore conformational space, particularly in the context of protein design [36, 56–58]. We hypothesized that by optimizing MSA cluster profiles with respect to AF2’s own confidence metrics, it may be possible to improve structure prediction without the need for experimental data. To test this hypothesis, we used ROCKET to optimize pLDDT, AF2’s primary confidence metric. This approach fails to improve the correspondence of AF2 predictions with experiment (Fig. S6), indicating that experimental data provide new and orthogonal information, beyond AF2’s implicit scoring function, that is necessary for efficient sampling of functionally relevant conformations.

### 2.4 ROCKET Overcomes Structural Barriers and Samples Diverse Conformations

We next asked whether ROCKET can perform challenging large-scale structural rearrangements – such as functional loop movements, peptide flips, and domain shifts. Such rearrangements are often beyond the scope of current state-of-the-art methods but may be easy to sample in MSA cluster profile space. Fig. 3 shows ROCKET-produced loop rearrangements in c-Abl kinase (Fig. 3a) and PTP-1B (Fig. 3b), as well as subunit rebuilding of *E. coli* GroEL (Fig. 3c). In all three cases, ROCKET successfully samples barrier-crossing transitions that are critical for modeling functional states. Beyond these large-scale motions, ROCKET also accurately refines domain rearrangements and smaller structural modifications, such as peptide flips in high-resolution structures (Fig. 3d-f). In contrast, PredictAndBuild, despite its own iterative use of AF2, largely fails to access these conformational changes (Fig. S7).

**Fig. 3.**
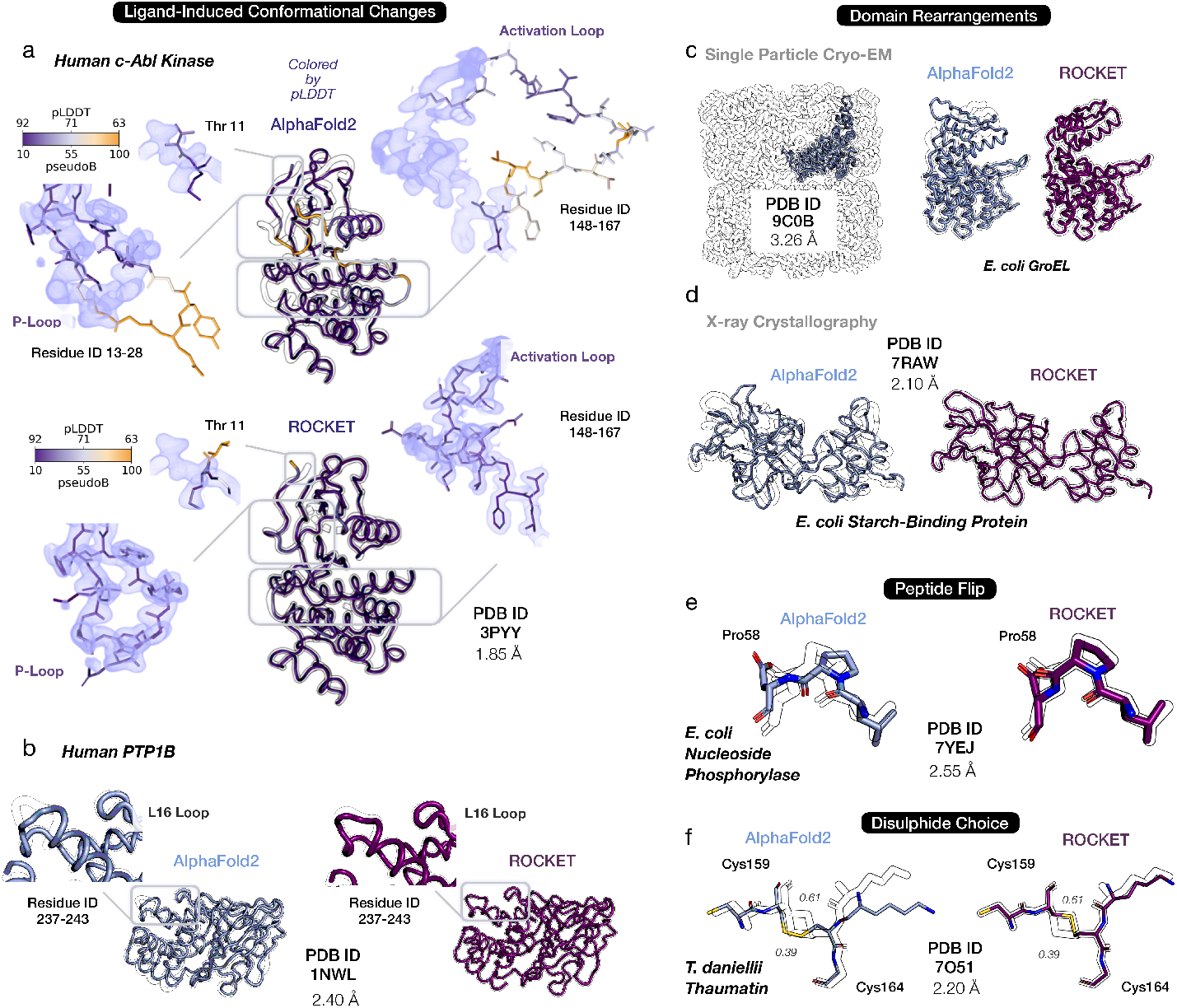
ROCKET Overcomes Structural Barriers and Samples Diverse Conformations. Shown are representative cases that highlight ROCKET’s ability to sample structural changes inaccessible to conventional refinement methods. In each panel, predictions by AF2 (top or left; light blue) and ROCKET (bottom or right; dark purple) are superimposed on experimental structures (black outline). Insets highlight local regions that undergo major changes. (a)Ligand-induced loop rearrangement in human c-Abl kinase (PDB ID 3PYY). ROCKET produces large conformational shifts of the activation and P-loops upon ligand binding. Per-residue confidence (pLDDT; blue-orange color scale) from AF2 is tracked by ROCKET and found to positively correlate with the final prediction’s agreement with experimental data. (b) Conformational changes in protein tyrosine phosphatase PTP-1B (PDB ID 1NWL). A peptide flip in the L16 loop, induced by ligand binding, is captured by ROCKET. Ligand molecules (not shown) are not considered by ROCKET, as its refinement depends purely on experimental density without explicit modeling of protein-ligand interactions. (c-d) Domain rearrangements for a cryo-EM dataset of *E. coli* GroEL (PDB ID 9C0B) and a crystallographic dataset of *E. coli* starch-binding protein (PDB ID 7RAW). (e) A peptide flip in the refinement of *E. coli* nucleoside phoshorylase (PDB ID 7YEJ). (f) A switch in a disulphide bond in the structure of thaumatin from *T. daniellii* (PDB ID 7AOJ). Two alternate conformations are present in the deposited structure, with refined occupancies of 0.39 and 0.61. ROCKET builds the conformation with highest occupancy.

Intriguingly, we find that, across our datasets, ROCKET’s AF2-derived model confidence (measured as per-residue pLDDT) is correlated with agreement with experimental data, as reflected in the positive correlation between final model confidence and RSCC (Fig. S8). We visually highlight an example of this in the refinement of the activation- and P-loops of c-Abl kinase (Fig. 3a). Since MSAs reflect the variety of conformations a protein can adopt [35], we suggest that ROCKET’s data-guided optimization uses, and then resolves, this structural ambiguity, allowing it to reach functional conformations that conventional refinement protocols cannot access. This clarifies, but does not contradict, our finding from the preceding section that pLDDT-driven optimization cannot refine predicted structures. Specifically, it indicates that pLDDT can identify highly preferred conformations but cannot, by itself, distinguish the experimentally observed state (and provide a useful gradient) without further experimental information.

### 2.5 ROCKET Integrates MSA Subsampling to Escape Local Minima in Large Transitions

The effectiveness of a gradient descent approach is inherently limited by local optima. When an initial AF2 prediction deviates significantly from the experimentally observed conformation, the gradient signal derived from the experimental likelihood target may become weak or misleading, impeding refinement. Inspired by recent findings that alternative folds can be uncovered through subsampling of the input MSA [35], ROCKET subsamples input MSAs to obtain diverse MSA cluster profiles – and thereby fresh initial models – that may lie closer to the experimental conformation. MSA subsampling in ROCKET follows the method outlined by AFCluster [35] and includes both clusters identified by DBSCAN and uniformally sampled MSAs. For each subsampled MSA, a prediction is made (as well as one from the full MSA) and scored using the experimental likelihood. The MSA resulting in the prediction with the highest likelihood (and thus most likely to accurately capture the experimentally resolved backbone arrangement) is subsequently selected for full refinement.

We illustrate this technique using the 1.8 Å crystallographic dataset of the human serpin PAI-1, captured in its energetically hyperstable latent state (Fig. 4a; left). The initial AF2 prediction, based on a full sequence alignment, adopts the protein’s metastable active conformation and deviates substantially from the experimentally determined model (Fig. 4a; right; 10.77 Å overall RMSD and 40.4 Å loop RMSD). A notable structural difference involves the reactive center loop (RCL, residues 330–354): it undergoes a large-scale structural transition from a solvent-exposed position in the active state to being fully inserted into the central *β*-sheet in the latent conformation. This rearrangement poses a substantial challenge for gradient descent optimization when refinement is started from the full MSA (Fig. 4b; 3.2 Å overall RMSD and 36.8 Å loop RMSD). On the other hand, subsampled MSAs result in starting conformations that more closely resemble the latent state (Fig. 4c; 4.2 Å overall RMSD and 13.6 Å loop RMSD for best starting structure). We note here that the experimental likelihood score does not monotonically correlate with pLDDT (Fig. 4c; lower panel), indicating that MSA selection based solely on AF2’s confidence metric offers no assurance of identifying the relevant functional state. ROCKET subsequently uses the highest likelihood MSA to perform gradient-based optimization, producing a structure that closely resembles the latent state (Fig. 4d; 1.9 Å overall and loop RMSD).

**Fig. 4.**
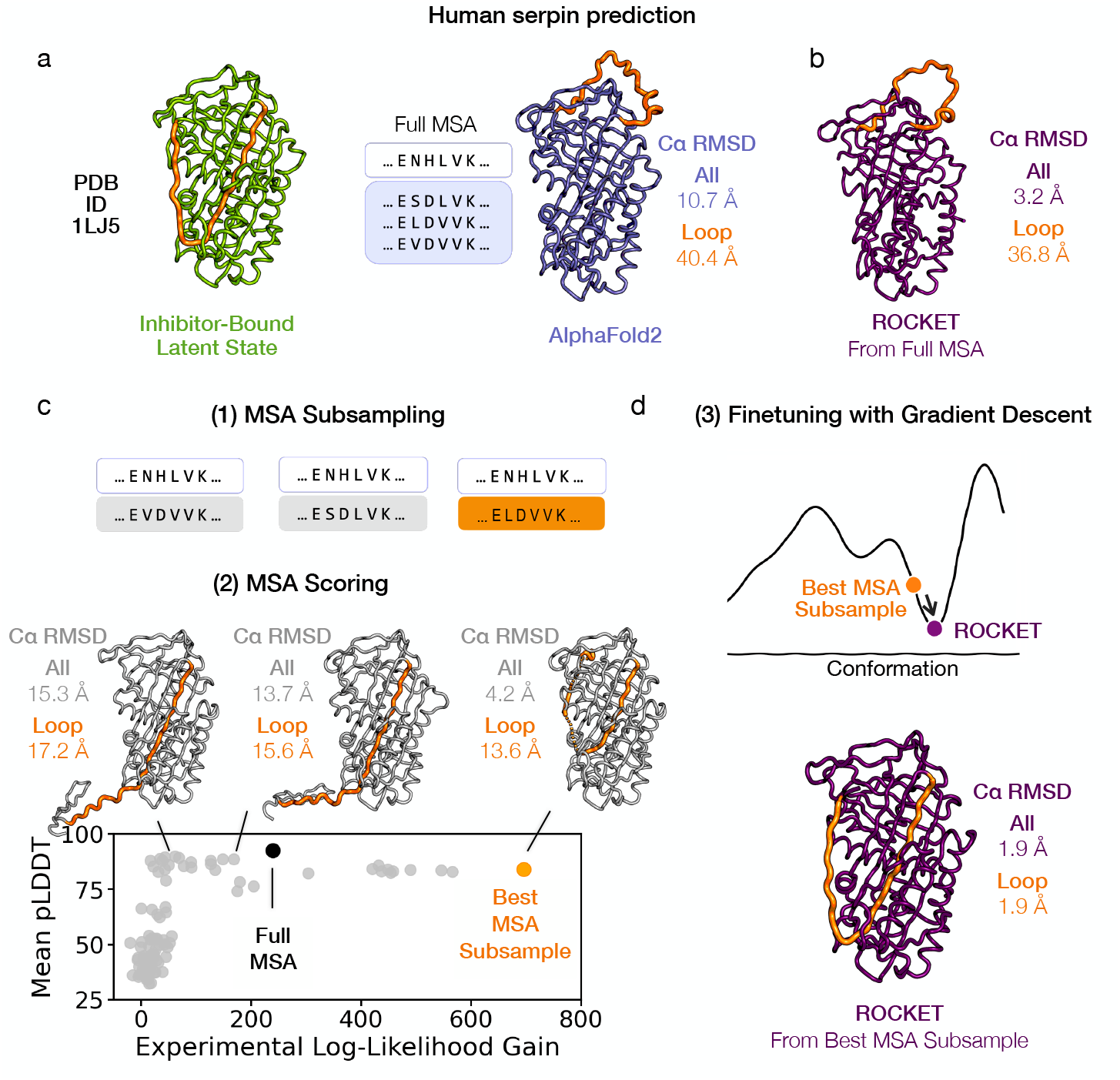
Synergistic Integration of MSA Subsampling with Gradient-Based Optimization. Gradient-based refinement faces a limit when initial predictions deviate substantially from the target conformation. (a) The structure of a human serpin, plasminogen activator inhibitor-1 (PAI-1), in its latent state (PDB ID 1LJ5, 1.8 Å) illustrates this issue (left). A conventional AF2 prediction, generated from the full MSA, yields the metastable active conformation that diverges from the latent state captured by the experimental structure (right). (b) The latent state conformation is not accessible by gradient descent alone when ROCKET refinement is started from the full MSA (c) Subsampled MSAs (1) result in predictions that more closely resemble the experimental conformation (2). Ranking these predictions (bottom panel) by their experimental likelihoods identifies a better starting model for gradient-based refinement (samples are ordered along the x-axis by increasing gains in experimental likelihood), unlike pLDDT-based scoring, which does not (highest samples along the y-axis do not resemble the experimental conformation). (d) Gradient-based refinement of the best starting prediction from (c) results in a structure that closely resembles the latent conformation.

### 2.6 ROCKET Extracts Essential Information from Low-Resolution Data

We next assess ROCKET’s applicability in the low-resolution regime with two datasets that lie past the typical resolution limits for automated model building: a 3.82 Å human HAI-1 crystallographic dataset and a 9.60 Å sub-tomogram average of *E. coli* GroEL.

In well-resolved regions of HAI-1, ROCKET successfully converges to a backbone conformation that more closely matches the experimental density than the AF2 prediction (Fig. 5a-b; regions 1, 2, and 5). Conversely, in areas where the experimental map is poorly resolved – such as region 3, which is not modeled in the deposited structure – ROCKET refrains from forcing arbitrary changes and instead preserves the original AF2 prediction. Particularly noteworthy is region 4 (residues 310–330), where the density is highly noisy, making even manual model building difficult. In this segment, where the deposited model appears to have an incorrect sequence register, ROCKET improves AF2’s initial prediction to better align with the density map without introducing new geometric outliers (Fig. 5c).

**Fig. 5.**
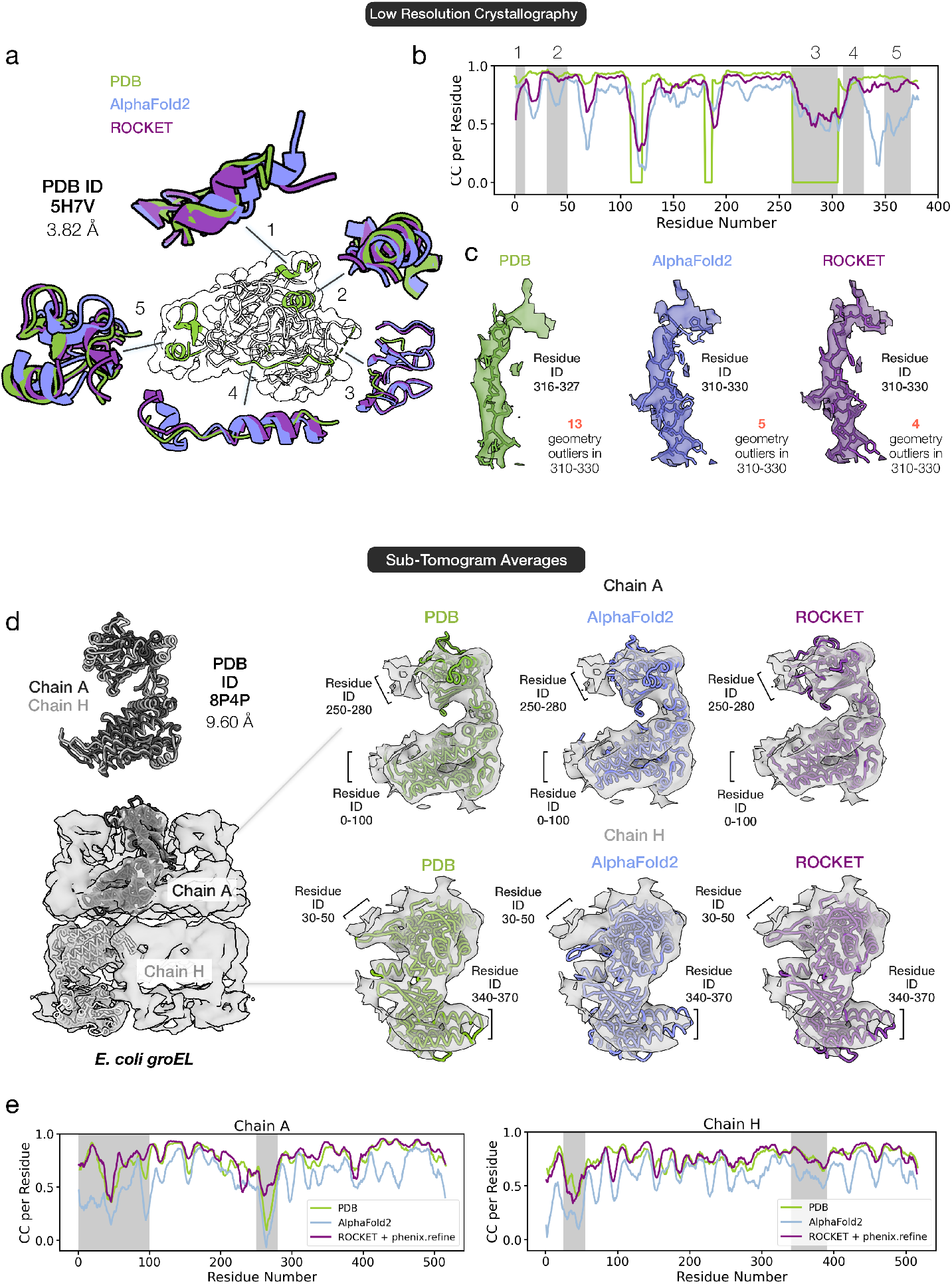
Extracting Information from Low-Resolution Data. (a-c) Refinement of a 3.82 Å crystallographic dataset of the human protease inhibitor HAI-1 (PDB ID 5H7V). ROCKET accurately recovers backbone structures in regions where the data support the deposited model (regions 1, 2, and 5) and preserves the AF2 prediction in poorly defined regions (region 3). In region 4, where manual model building is hindered by noisy density and seemingly incorrect sequence register in the deposited model, ROCKET improves the initial AF2 prediction without introducing new geometric outliers. (d-e) Refinement of a 9.60 Å sub-tomogram average of *E. coli* GroEL (PDB ID 8P4P). ROCKET successfully predicts two distinct subunit conformations observed in the GroEL heptameric rings. Maps computed using the refined models correlate with the experimental map at a level comparable to human-built models. For the chain A conformer, ROCKET explores a broader conformational space, with its final model achieving a higher average RSCC of 0.5 for residues 250–280 (marked by a star) than the deposited model’s RSCC of 0.2.

In the sub-tomogram average of GroEL (PDB ID 8P4P), the 9.60 Å map reveals two distinct conformations for the repeating subunit in the top and bottom heptameric rings (Fig. 5d). Despite this very low resolution, ROCKET accurately predicts both conformations (Fig. 5d), yielding refined models with map correlations comparable to manually built structures (Fig. 5e). Moreover, for chain A, ROCKET explores a broad conformational space, deviating substantially from the AF2 prediction. Notably, in the top domain (residues 250–280; marked by a star in Fig. 5e), ROCKET achieves an average RSCC of 0.5, substantially higher than the 0.2 achieved by the deposited model, supporting a closer match to experimental data.

These examples demonstrate that ROCKET extracts meaningful information at resolutions inaccessible to other methods. Moreover, in challenging regions such as the 310–330 helix of HAI-1 or the 250–280 residue range of GroEL chain A, ROCKET produces solutions that improve agreement with the experimental map while retaining structural plausibility, surpassing the accuracy of human-deposited models.

## 3 Discussion

In developing ROCKET, we have demonstrated that experimentally guided refinement of the embedded MSA profiles of AF2 enables efficient exploration of conformational space, capturing structural changes that conventional refinement in Cartesian space struggles to reach. Our results suggest that these embeddings provide access to paths along which the barrier for structural rearrangements is significantly reduced or eliminated, indicating that information about such rearrangements are encoded in evolutionary statistics.

We anticipate that data-guided, inference-time optimization will prove broadly valuable across diverse atomic model building scenarios. Beyond cryo-EM and crystallographic datasets, implemented here, our approach can be used for other, sparser data modalities, provided a likelihood-based target between data and prediction can be formulated. Extending ROCKET to handle multi-chain complexes or protein-ligand co-folding is straightforward within any AlphaFold-like framework, requiring in principle only a switch in the inference model used.

More ambitious directions may involve integrating generative models to account for conformational ensembles. Futher, given our success with MSA profile biasing, we propose that learning a mapping from experimental observables to a profile bias matrix could effectively condition single-shot structure prediction towards experimentally probed conformations. Such a mapping would translate experimental data – such as cryo-EM density maps, X-ray diffraction intensities, or NMR-derived constraints – into a profile bias matrix that captures residue-level probabilities or pairwise constraints derived from experiment, effectively guiding the model towards relevant conformations without requiring exhaustive searches. By adopting amortized inference, which learns a reusable mapping to enable fast predictions for new inputs, we could further streamline the process by replacing the stepwise search (MSA subsampling and likelihood scoring, shown in Fig. 4) with a more efficient learned transformation.

While additional work is needed to fully realize these extensions, our study marks critical progress by showing that structure optimization in coevolutionary embeddings can overcome limitations of conventional refinement. Additionally, by introducing differentiable likelihood targets for cryo-EM and crystallography, which include a robust treatment of measurement error, we provide a framework well-suited for training future predictive models. More broadly, our Bayesian approach to combining information learned by ML models with information obtained by direct observation establishes a foundation for continuous interaction between machine learning and experimental data. This interplay is critical for the interconnected goals of scaling experimental throughput and training machine learning models with enhanced functionality.

Our atomic model building is automated, requires no retraining, and for difficult cases produces models of quality comparable or even superior to those created by expert human modelers. However, certain limitations remain. The current approximation of the atomic displacement parameters is derived empirically from model confidence, and could be improved by incorporating considerations of density fit at the residue level. Additionally, because OpenFold is not explicitly aware of crystal contacts, ROCKET may struggle to converge to certain lattice-constrained conformations. We anticipate that this limitation can be addressed by extending ROCKET to use multi-chain models [59]. We also noticed that ROCKET can fail to flip small loops (3–4 residues in length) that contain bulky sidechains. We show examples of cases that are difficult for ROCKET in Fig. S9. Perhaps most importantly, iterative backpropagation through OpenFold is memory-intensive and limits the maximum size of the protein or domain that can be refined at once – about 500 residues on a 40 GB A100 GPU (Fig. S10). We believe this can be extended with further code optimization; the implementation of a learned mapping for single-shot MSA profile bias, mentioned above, could also help address this limit.

Naturally, assessing biological accuracy solely through the lens of how well the atomic model fits experimental maps has inherent limitations, as both cryo-EM and crystallography can introduce artifacts [60]. Ultimately, multi-modal information is essential for building a full picture of physiological protein states and functions. For this purpose, our approach is readily adaptable to alternative loss functions that combine multiple sources of experimental and computational data, supporting integrative modeling strategies for biological structure determination.

## 4 Methods

### ROCKET Algorithm and Processing Pipeline

ROCKET’s inputs and pre-processing steps are summarized in Algorithm 1, while the ROCKET inference-time optimization algorithm is summarized in Algorithm 2.

### Inputs and Pre-Processing Pipeline

For crystallographic datasets, a protein sequence and a reflection MTZ file (or CIF file) containing observed intensities and their uncertainties are required, while for cryo-EM datasets two half-maps are required. As outlined in Algorithm 1, to obtain an aligned reference model (**x**^ref^), we use Phasertng [61] for molecular replacement with crystallographic datasets and a likelihood-based docking tool, EM placement [62], with cryo-EM datasets. These determine the correct pose from an initial, unconditional AF2 model. These tools also estimate experimental data parameters for refinement, such as *E*_*e*_ and *D*_*obs*_, which represent observed normalized amplitudes and an accounting for measurement error, respectively [63], and are described further below.

#### Algorithm 1

ROCKET Preprocessing

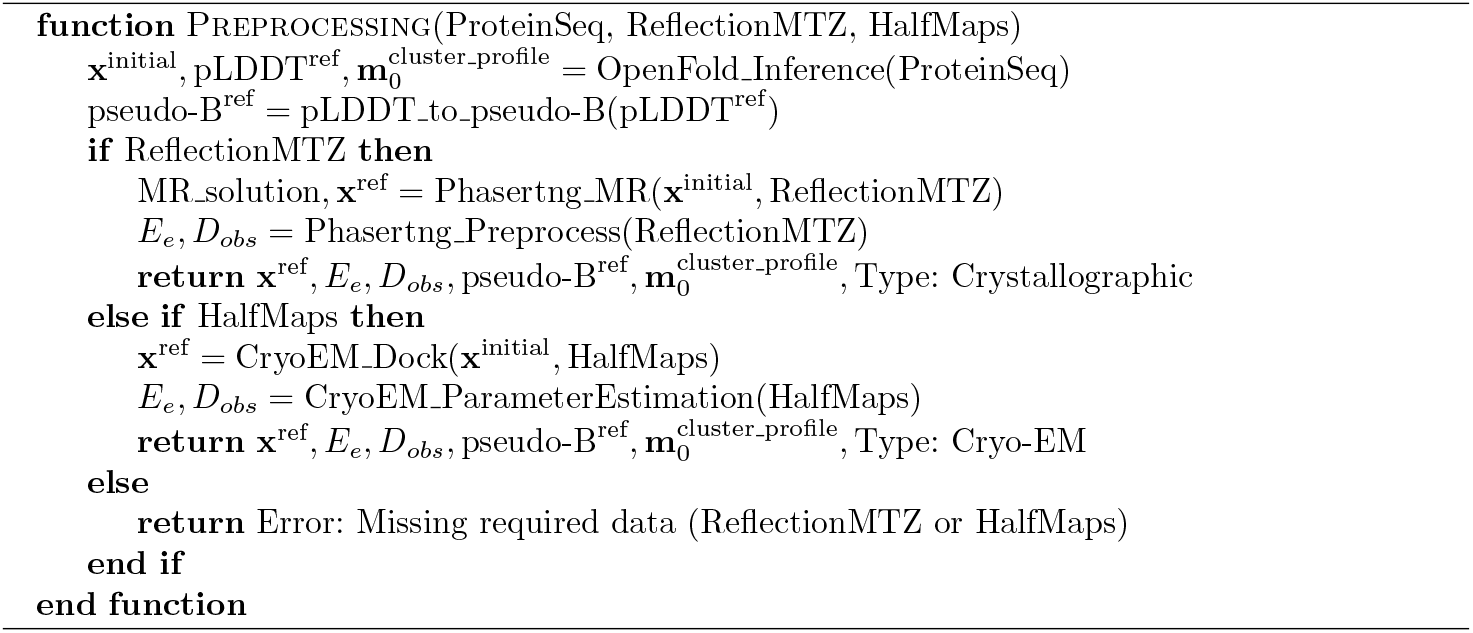

### Refinement Algorithm

ROCKET optimizes a linear bias (with scales **w** and offsets **b**) that it applies to the starting MSA cluster profile 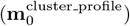 to maximize agreement between an OpenFold prediction (**x**^prediction^) and experimental data (*E*_*e*_). This agreement is quantified by a data log-likelihood gain (LLG) target, *L*_LLG_, which is combined with an optional positional restraint, *L* _L2_, to yield an overall objective function *L* (see below). The shapes of the **w** and **b** tensors match that of the MSA cluster profile tensor ([number of MSA clusters x number of residues x 23] [1])

During each ROCKET iteration, we apply a linear bias to the initial cluster profile matrix and then run OpenFold inference to obtain a new prediction, along with its pLDDT confidence values. Our current implementation estimates atomic B-factors at every iteration from pLDDT confidence scores using a previously established heuristic [2], without explicitly refining them. Specifically, we convert

#### Algorithm 2

ROCKET Refinement

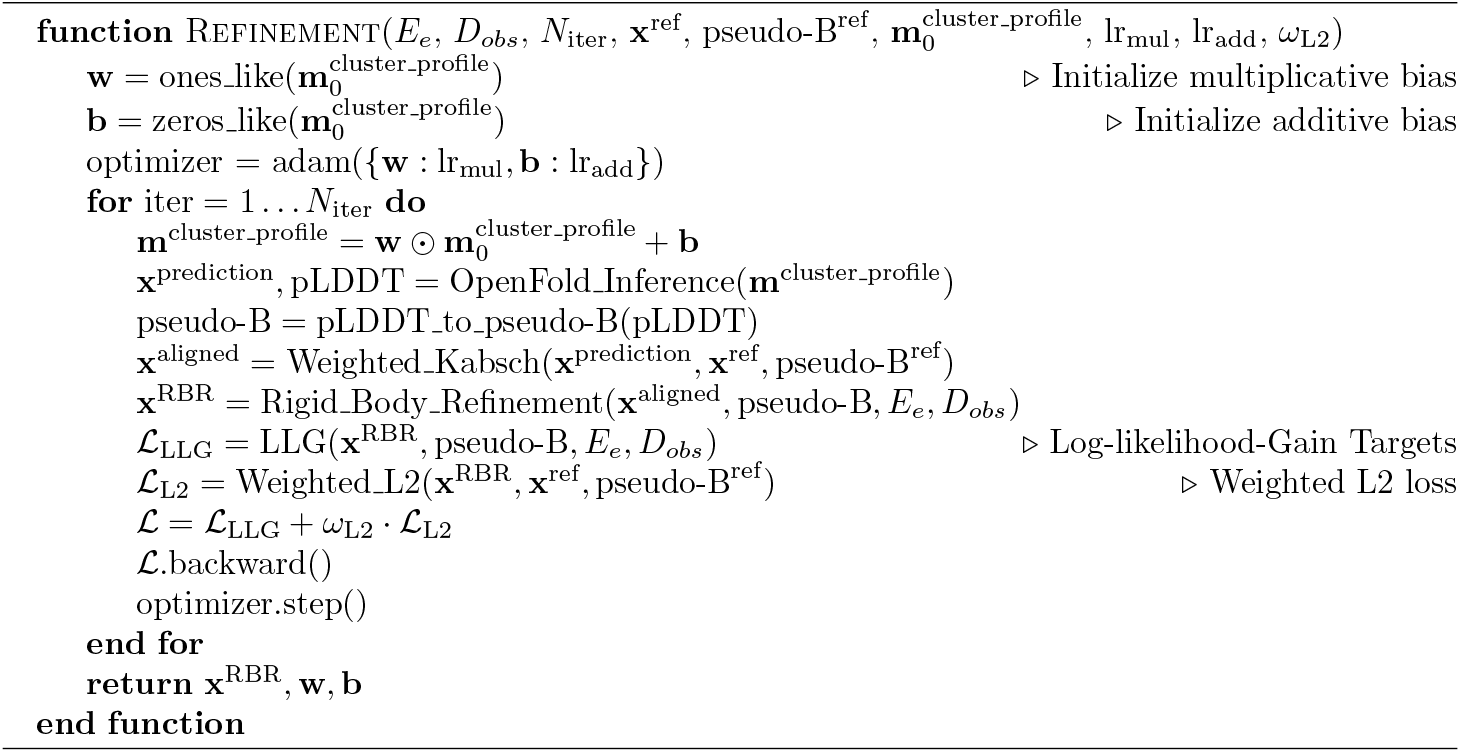

pLDDT values into equivalent RMSD values, using the empirical relation from Baek et al. [2], and then to corresponding pseudo B-factors:

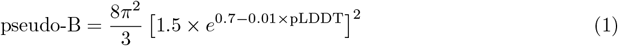

At every iteration, we align the newly predicted model to the reference model using a weighted Kabsch alignment (with pseudo B-factors serving as the weights), followed by rigid-body refinement (elaborated in a separate section below). We then compute the experimental log-likelihood-gain (LLG) using the aligned coordinates and the pseudo B-factors.

In practice, ROCKET runs in two phases: an “adventurous” phase 1 and a “fine-tuning” phase 2. We apply different learning rates for the multiplicative (**w**) and additive (**b**) components of our MSA profile bias, and these rates vary across phases. In phase 1, we use higher learning rates (by default, lr_mul_ = 1.0 and lr_add_ = 0.05) and a default low-resolution cutoff of 3 Å. We also incorporate a weighted *C*_*α*_ mean squared distance between the reference model and latest prediction as an L2 regularization term that quantifies initial model confidence. The weights are computed using the same scheme we employ for the weighted Kabsch alignment discussed further below:

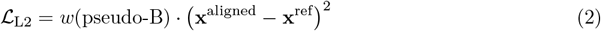

with the confidence-based weights *w*(pseudo-B) defined in Equation 13 and the final loss for backpropagation:

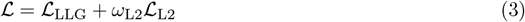

By default, the L2 loss weight is *ω*_L2_ = 10^*−*11^. We run phase 1 for three independent traces, each consisting of 100 iterations, and select the model with the best LLG score to proceed to phase 2. The aim of phase 2 is to further fine-tune the structure. By default, we set phase 2 to run for 500 iterations with both lr_mul_ and lr_add_ to 10^*−*3^ and remove the L2 loss term (*ω*_L2_ = 0). An early stop occurs if the LLG score does not improve by more than 0.1 for 50 consecutive iterations. Figure S12 illustrates the efficacy of phase 1 for avoiding local optima through the example of the c-Abl kinase crystallographic dataset (PDB ID 3PYY).

The pseudo-B approximation can limit accuracy by not capturing finer structural details. Moreover, geometric validation indicates that outputs from the iterative optimization have more bond outliers and steric clashes than stricter refinement protocols typically allow (Fig. S3a). To address these limitations, we append a short standard local-refinement step using phenix.refine [16] after iterative OpenFold inference. Analogous to AMBER relaxation in standard AF2 pipelines [1], this step further optimizes geometry (Fig. S3) and displacement parameters, polishing the final model’s overall quality.

### Implementation

ROCKET is implemented in PyTorch 1.12.1 as an extension of the OpenFold system. It currently uses the monomer version of OpenFold with AF2 model 1 weights to maintain consistency with AF2’s data splits. This allows for prediction and refinement of crystallographic datasets containing a single chain in the asymmetric unit or one domain (at a time) in a cryo-EM complex.

### Crystallographic Log-Likelihood-Gain Targets

For crystallographic datasets, we employ the log-likelihood-gain on intensity (LLGI) target introduced in [63], where for acentric and centric reflections:

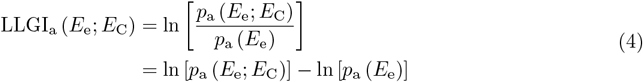

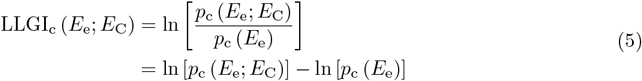

with 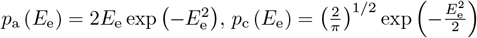, and:

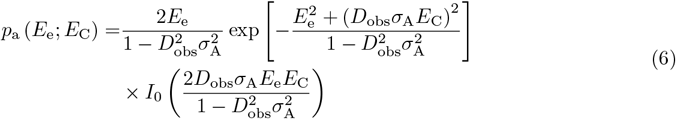

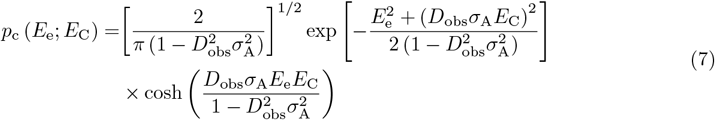

and *p*(*x*; *y*) denotes the conditional probability of *x* given *y*.

As defined in [63], *E*_*e*_ is the “effective” observed normalized amplitude, *E*_*C*_ is the normalized structure factor amplitude calculated from the predicted model in a differentiable manner using SFCalculator with solvent correction [34], *D*_*obs*_ encodes the reduction in correlation between true and “effective” normalized structure factors arising from experimental error, and *σ*_*A*_ is a resolution-dependent factor that encodes the reduction in correlation between the true and calculated normalized structure factors arising from model error.

We refine *σ*_*A*_ in resolution bins [64] at every ROCKET iteration using the Newton-Raphson optimization method [65]. The log-likelihood gain of the observed effective amplitudes, *E*_e_, given the calculated amplitudes, *E*_C_, is maximized by refining *σ*_*A*_. The derivative of the log-likelihood gain with respect to *σ*_*A*_ for each resolution bin is given by:

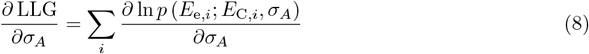

where *i* is an index over observations. The second derivative is given by:

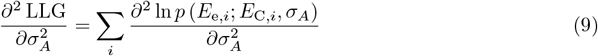

We obtain the updated value of *σ*_*A*_ using a Newton step:

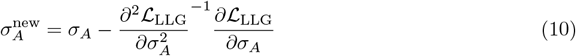

The value of *σ*_*A*_ is constrained within [0.015, 0.99] to maintain physical relevance and stability during refinement. For all LLG and *σ*_*A*_ estimates we use a working set of reflections (we find that using the working set for *σ*_*A*_ refinement does not lead to any meaningful overfitting with ROCKET, Fig. S11). We keep a test set of reflections for final R-free calculation.

### Log-Likelihood Gain Target and Noise Modeling for Cryo-EM

For cryo-EM data, we follow the method outlined in [62] to dock the initial prediction into the experimental map. We model the signal and noise in Fourier space to account for directional and resolution-dependent variations in data quality. The signal is derived from correlations between Fourier terms of the experimental half-maps, while the noise is deduced from their differences [52].

For a single Fourier term, the log-likelihood gain (LLG) is given by:

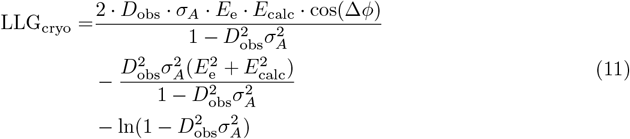

where *E*_e_ and *E*_calc_ are the normalized amplitudes of the observed and calculated Fourier terms, Δ*ϕ* = *ϕ*_calc_ *− ϕ*_obs_ is the phase difference between these Fourier terms, and *D*_obs_ and *σ*_*A*_ are analogous to their crystallographic counterparts.

We compute *σ*_*A*_ for each resolution bin as in [52], using observed normalized amplitudes (*E*_e_), calculated normalized amplitudes (*E*_calc_), and the phase difference Δ*ϕ* = *ϕ*_calc_ *− ϕ*_obs_.

### Weighted Kabsch Alignment and Rigid Body Refinement

As stated above, we use the initial prediction from OpenFold to run Phasertng (or EM placement for cryo-EM) for molecular replacement. For every iteration, we align the OpenFold model to the reference molecular replacement solution before computing the relevant LLG score. This alignment is performed by first solving the following minimization problem with the Kabsch algorithm [66]:

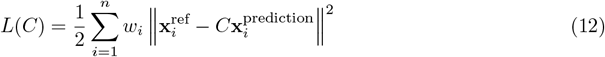

where *C* denotes the rotation-translation matrix, **x**^ref^ and 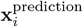 are corresponding atomic coordinates of atom *i* in the reference and moving model, respectively, and *w*_*i*_ are positional weights. Only *α*-carbon (*C*_*α*_) atoms are included in the alignment, and their weights are determined empirically from the pseudo-B values of the reference structure (the first, unconditioned OpenFold prediction). Specifically, for each residue:

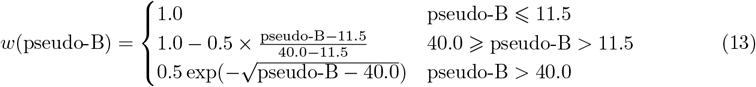

Translation vectors are determined by the vector difference of weighted center of mass of *C*_*α*_ atoms in the reference and moving models, then the rotation matrices are estimated with Kabsch algorithm. Once the alignment is completed, a subsequent rigid-body refinement is performed through gradient optimization of the LLG target:

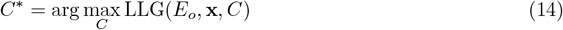

where **x** is the model coordinates after Kabsch alignment. When the predicted aligned error (PAE) matrix from AF2 can be used automatically to split proteins into domains, ROCKET performs domain-specific alignments to make the best use of gradient information from the LLG. We used this, for example, for the refinement of the dataset related to PDB ID 7RAW in Fig. 3c.

### Multi-Chain Dataset Handling

ROCKET can readily handle monomeric protein predictions in its current form. We have also demonstrated refinement of crystallographic datasets that contain two chains in the asymmetric unit for the kinase datasets (PDB ID 3PYY in Fig. 3 and PDB ID 7DT2 in Fig. S9). For these cases, we included the second chain present in the asymmetric unit as a fixed Fourier contribution in the likelihood calculation but excluded it from refinement. However, general multi-chain refinement would require integrating ROCKET with a multimeric model such as OpenFold-Multimer.

Given the current size limit of about 500 residues (on 40GB GPUs) for backpropagating through OpenFold, we also wanted to illustrate how larger cryo-EM complexes may be refined in parts. For the groEL example in Fig. 3c (PDB ID 9C0B), we only refined the domain of the groEL subunit that requires the most rebuilding. We did so by inserting a 9-glycine loop in the region connecting the two domains of the subunit, and treating the resulting single entity as the target for prediction.

## Data Visualization

Visualization of PDB structures and experimental densities was done with PyMOL [67] and Moorhen [68].

## Code Availability

Release of all ROCKET code and target functions is in preparation.

## Acknowledgments

The following funding is acknowledged: Wellcome Trust (grant No. 209407/Z/17/Z to RJR); National Institutes of Health (NIH), National Institute of General Medical Sciences (P01-GM063210 to RJR and TCT; R35-GM150546 to MA; and DP2-GM141000 to DRH); Biotechnology and Biological Sciences Research Council (grant No. BB/Y009398/1 to RJR); a graduate fellowship, Eric and Wendy Schmidt Center to MHL. We would also like to thank K.M. Dalton, R. Gaudet, T.J. Lane, S. Eddy, and S. Petti for insight and discussions, and C. Floristean for help with the OpenFold codebase.

## Supplementary Table

**Table 1.** Table of high-resolution crystallographic benchmark dataset used for ROCKET. n_res indicates the number of residues.

| Code | n_res | TM_score | Spacegroup | Resolution (Å) | RMSD(AF2, deposited) (Å) |
| --- | --- | --- | --- | --- | --- |
| 7E3Z | 261 | 0.974 | P 1 21 1 | 1.45 | 1.29 |
| 7VNX | 223 | 0.996 | P 61 | 1.80 | 0.75 |
| 7DNT | 250 | 0.990 | P 21 21 2 | 2.50 | 1.74 |
| 7O51 | 207 | 0.994 | P 41 21 2 | 2.20 | 0.90 |
| 7TFQ | 292 | 1.000 | P 21 21 2 | 1.75 | 0.48 |
| 7RAW | 251 | 0.975 | I 21 3 | 2.10 | 0.89 |
| 7FIU | 293 | 0.943 | P 21 21 21 | 1.84 | 1.77 |
| 7SEZ | 221 | 0.990 | C 2 2 21 | 1.70 | 1.75 |
| 7RM7 | 228 | 0.938 | P 21 21 21 | 1.03 | 0.54 |
| 7QDV | 100 | 0.994 | P 43 21 2 | 1.90 | 0.84 |
| 7TRW | 217 | 0.994 | P 62 | 2.28 | 1.96 |
| 7T26 | 144 | 0.999 | P 41 21 2 | 1.14 | 0.67 |
| 7S3L | 269 | 0.997 | P 43 21 2 | 2.60 | 1.56 |
| 7DMS | 94 | 0.424 | P 21 21 21 | 1.96 | 0.56 |
| 7AOJ | 179 | 0.910 | P 61 | 1.63 | 1.60 |
| 7U2R | 245 | 0.999 | P 61 2 2 | 1.85 | 0.54 |
| 7KZH | 202 | 0.979 | P 3 2 1 | 2.49 | 0.62 |
| 7EJG | 237 | 0.998 | P 61 | 1.68 | 1.11 |
| 7T7Y | 154 | 0.990 | P 2 21 21 | 1.81 | 0.68 |
| 7EDC | 249 | 0.989 | C 1 2 1 | 1.95 | 1.46 |
| 7EYJ | 95 | 0.999 | P 62 | 1.38 | 0.64 |
| 7RPY | 256 | 0.957 | P 32 2 1 | 1.67 | 0.84 |
| 7ECD | 272 | 0.997 | I 41 | 2.60 | 1.31 |
| 7UNN | 260 | 0.902 | P 21 21 21 | 1.45 | 0.58 |
| 7TBS | 238 | 0.945 | I 2 2 2 | 1.96 | 0.62 |

## Supplementary Figures

**Fig. S1.**
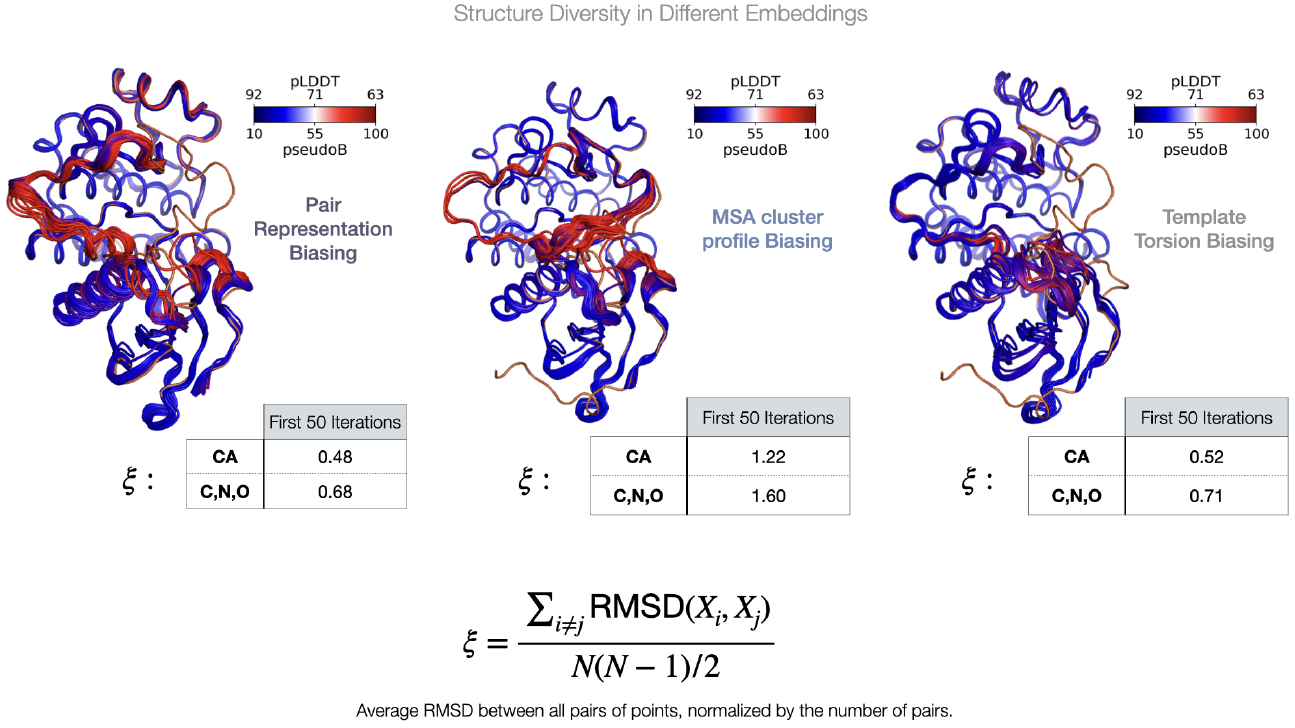
ROCKET Optimization in Different AlphaFold2 Embeddings. We performed data-guided inference-time optimization, as described in the main text, in alternative embeddings. We found that optimizing MSA cluster profiles provides the highest structural diversity along search trajectories. We calculated an average RMSD (*ξ*) between all pairs of structures, normalized by the number of pairs for the first 50 structures in a ROCKET phase 1 refinement run. Diversity is much higher when biasing the MSA cluster profile (*ξ* = 1.22 versus *ξ* = 0.48 and *ξ* = 0.52 when biasing the pair and template torsion representations, respectively).

**Fig. S2.**
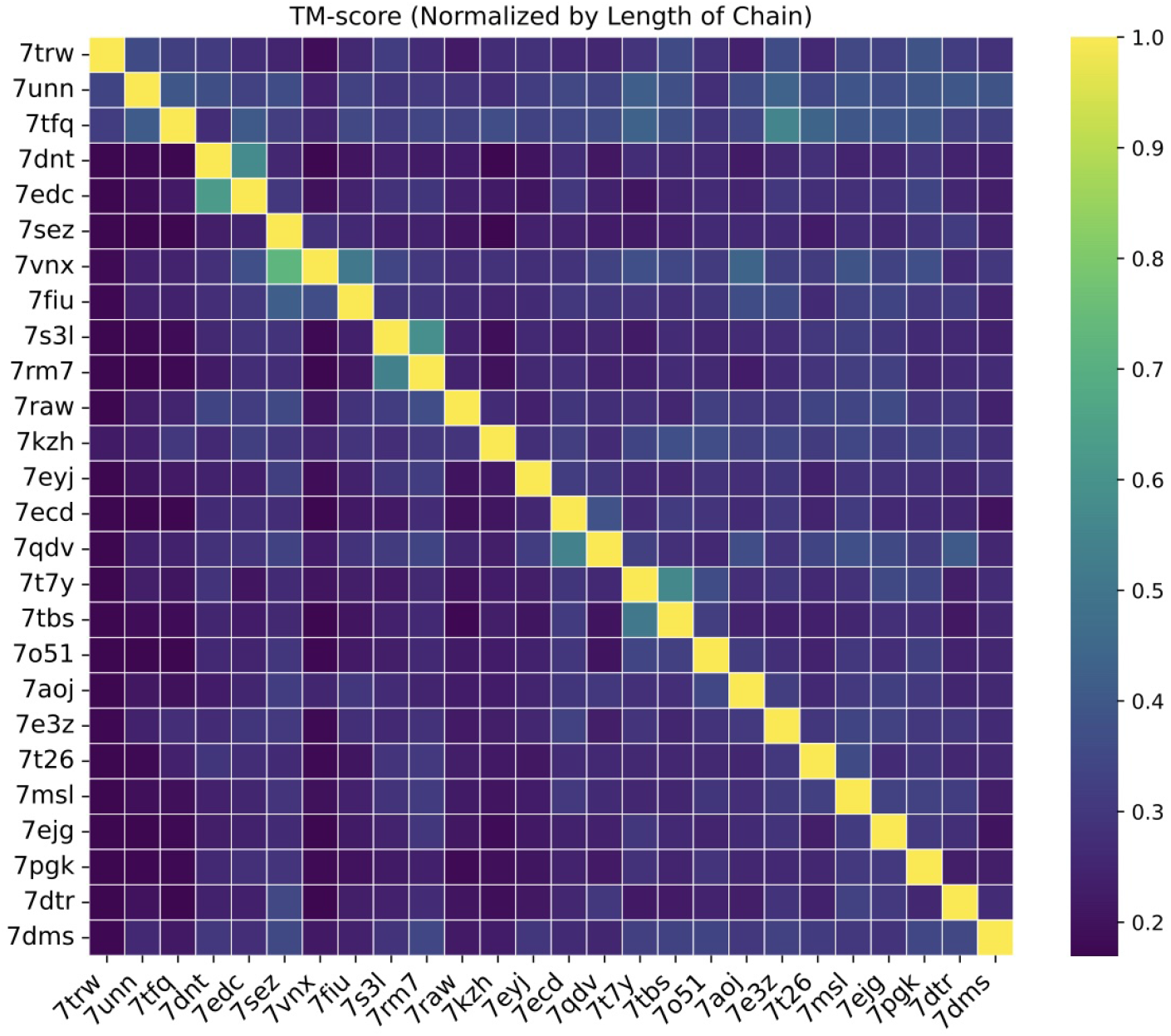
Fold Diversity in High-Resolution Benchmark Dataset. Template Modeling (TM) scores for the 27 structures present in our crystallographic benchmark dataset, identified through their PDB ID. The TM-score evaluates how well two structures align in three-dimensional space, independent of sequence identity, with TM-score closer to 1 indicating very high similarity in the two folds. The low TM-scores between structures reflect their fold diversity. These structures were all released after the training of the AF2 weights used here and were solved by the single-wavelength anomalous diffraction (SAD) method, suggesting that it was challenging to find structural homologs in the PDB for molecular replacement. We therefore expect limited leakage from the original AF2 training set.

**Fig. S3.**
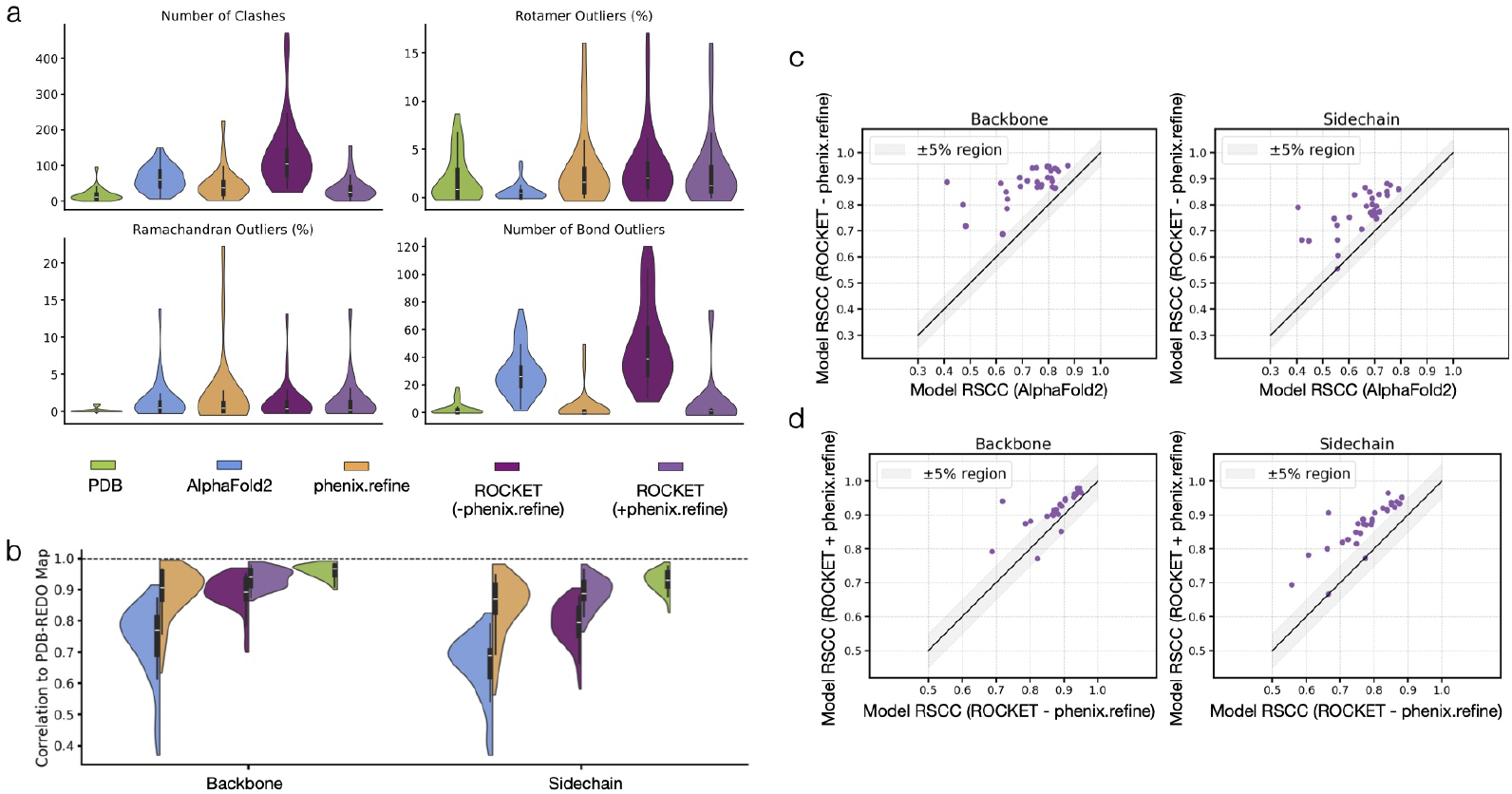
Refinement Results for High-Resolution Crystallographic Datasets. (a) Geometric validation after refinement by different methods, carried out using MolProbity [69] for the 27 high-resolution crystallographic test cases. (b)Real-space Pearson correlation coefficient (RSCC) values for backbone and sidechains across initial AF2 models, phenix.refine, ROCKET refinements without or with the phenix.refine step. All residues were included, including those for which AF2 and ROCKET report low confidence. RSCC values for human-deposited models are also shown. Note that PDB-REDO maps are expected to have favorable phase bias for these models, which will artificially increase reported RSCC. (c-d) Breakdown of the incremental RSCC improvement from the initial AF2 prediction to the ROCKET structures (c), and before and after the phenix.refine step (d).

**Fig. S4.**
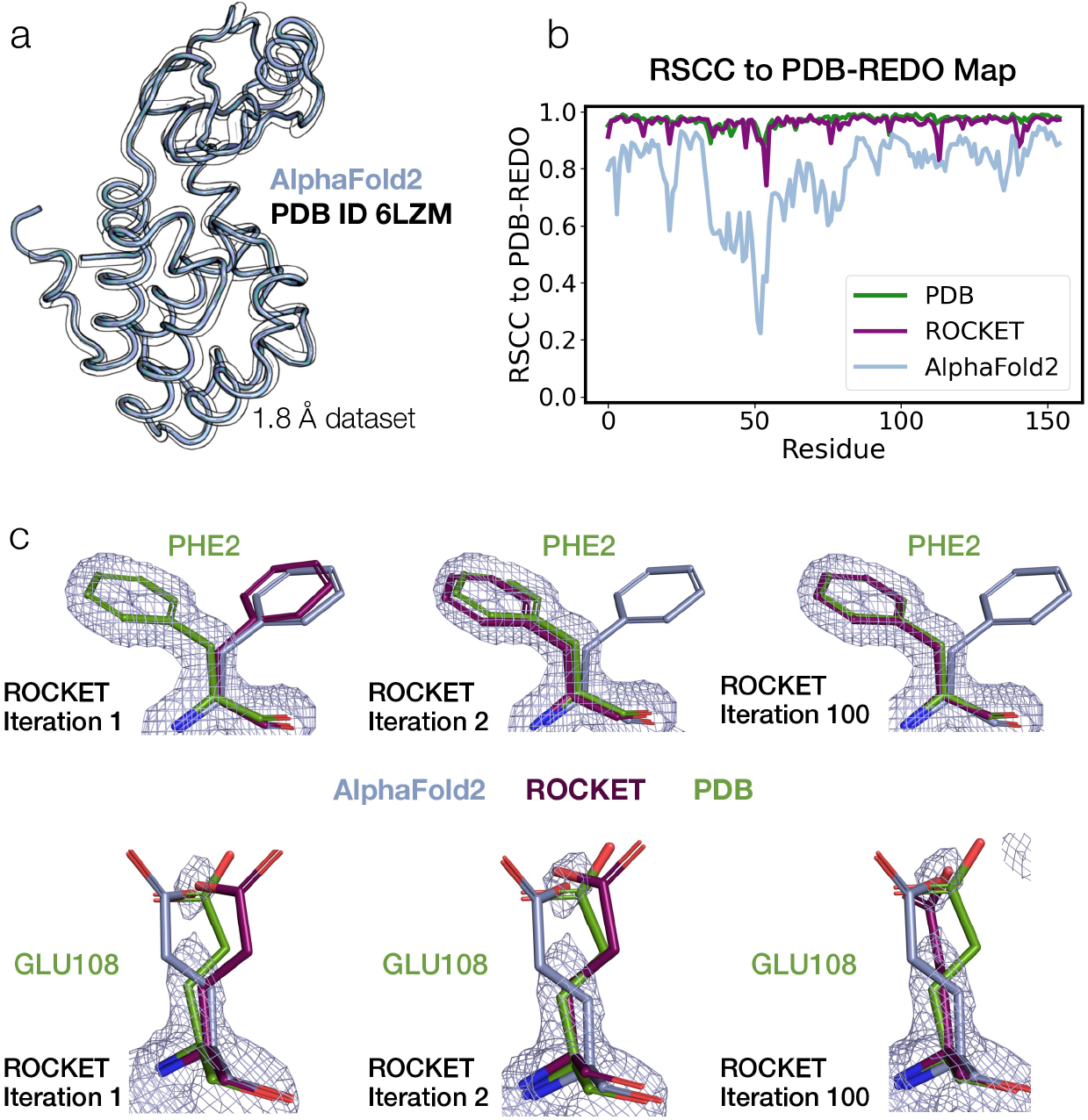
MSA Cluster Profile Fine-Tuning Enables Both Backbone and Sidechain Refinement. Example of ROCKET refinement for a bacteriophage T4 lysozyme dataset (related PDB ID 6LZM), which differs from the AF2 prediction, particularly in the orientation of its upper domain (a). The PDB REDO structure is shown in transparent outlines. (b) ROCKET can refine the prediction to the same quality as the deposited structure. (c) When the data support a clear conformation (*e*.*g*., for PHE2), ROCKET is able to find different rotameric configurations and gradually fine-tune the sidechain position. For GLU108, where the data are noisy, ROCKET explores different conformations and, in this case, settles for placing a carboxylate oxygen atom in the available density.

**Fig. S5.**
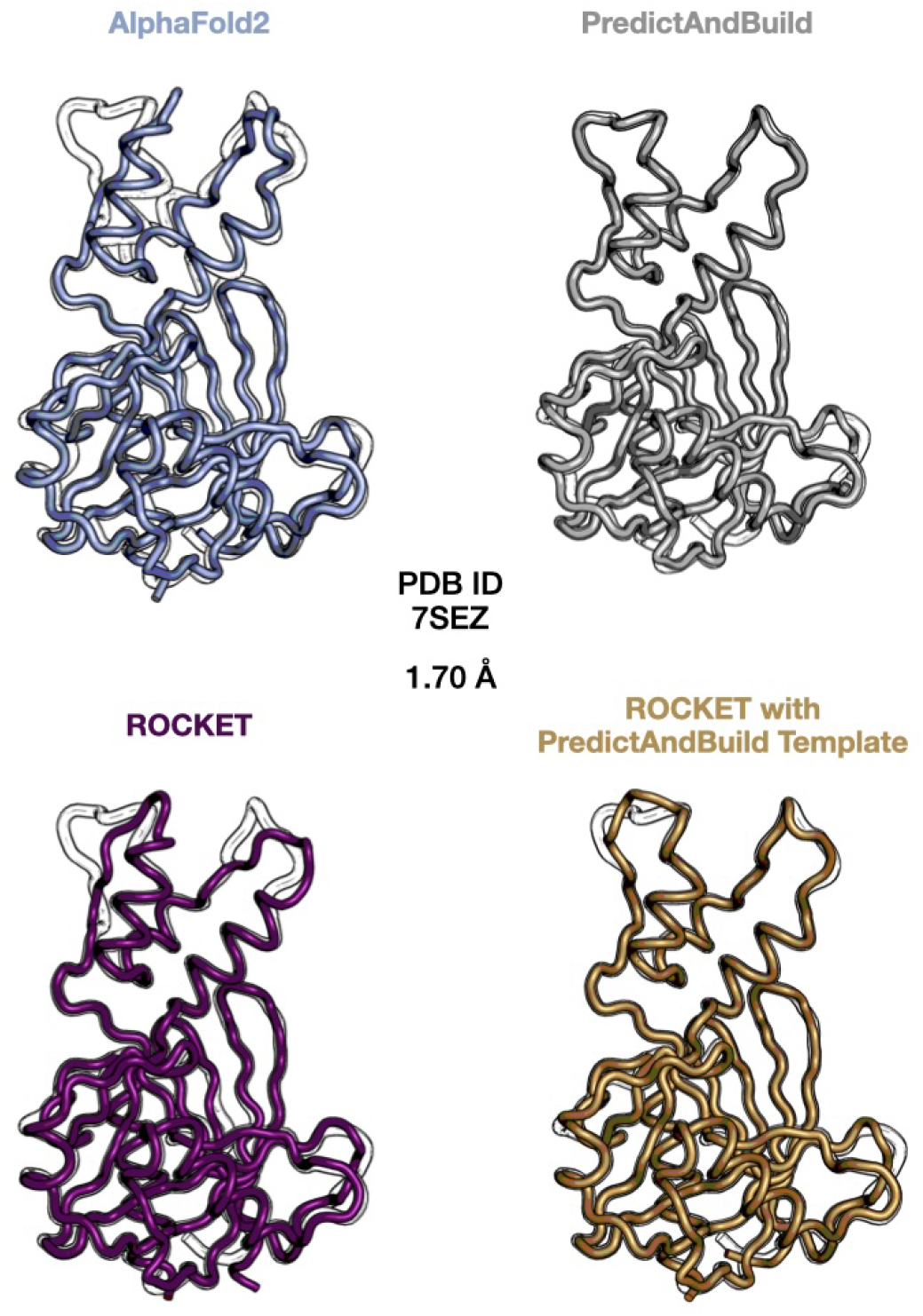
Combining ROCKET with PredictAndBuild. We discuss in the main text and further in Fig. S9 that ROCKET can struggle to flip small loops containing long sidechain residues. The refinement of the Vaccinia Virus decapping enzyme D9 (PDB ID 7SEZ) is an example of this, where ROCKET is unsuccessful in modeling two small loops in the upper domain, while PredictAndBuild converges to the correct backbone. We show the complementarity of the two approaches by running ROCKET with the PredictAndBuild structure provided as a template during OpenFold inference and improving the final output.

**Fig. S6.**
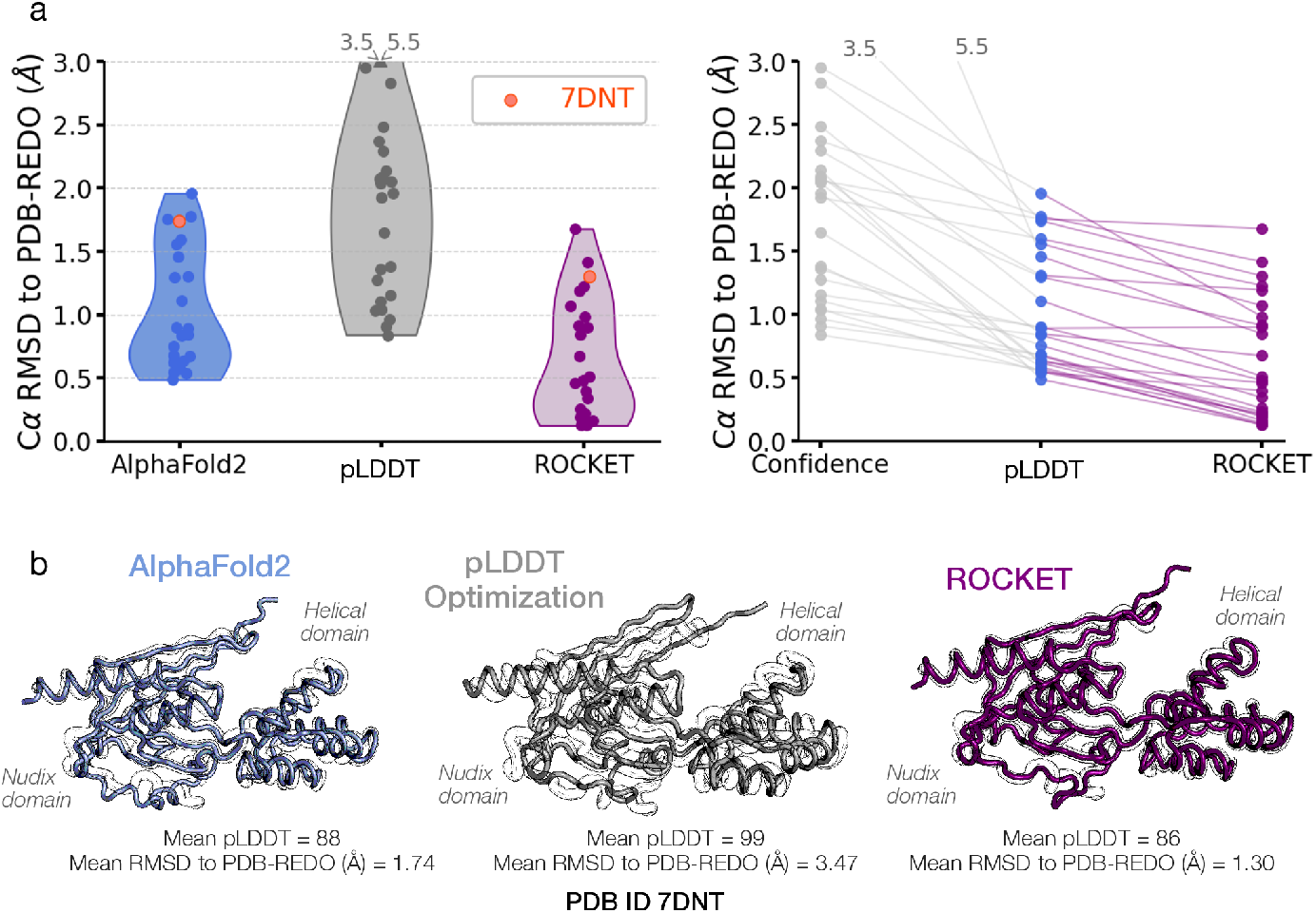
Comparison with Confidence Maximization. To assess the added value of experimental data integration, we compare ROCKET’s inference-time optimization to the results that can be obtained by maximizing AF2 model confidence. AF2 model confidence maximization has previously been used to explore alternate conformations for a given sequence, especially in the context of protein design. Depending on the task, different studies have optimized AF2’s local confidence metric (predicted local-distance difference test – pLDDT) [56–58], its confidence of relative positioning between residues (predicted aligned error matrix – PAE matrix) [56], or interface confidence for complexes [36, 58]. We conducted a search in the MSA cluster profile space that maximizes pLDDT and found that AF2 does not easily produce experimentally observed conformations without further experimental information. (a) C*α* RMSD values between PDB-REDO models and structures from standard AF2 inference, pLDDT maximization, and ROCKET’s data-likelihood maximization. ROCKET consistently improves the match between AF2 predictions and experimental structures, while pLDDT maximization alone does not achieve comparable accuracy. (b) Example of the viral mRNA-decapping enzyme g5rp (PDB ID 7DNT) illustrating ROCKET’s refinement capability. The initial AF2 prediction shows domain misalignment and secondary structure inaccuracies compared to the experimentally resolved conformation. ROCKET refines these discrepancies, while pLDDT maximization does not converge to the correct structure.

**Fig. S7.**
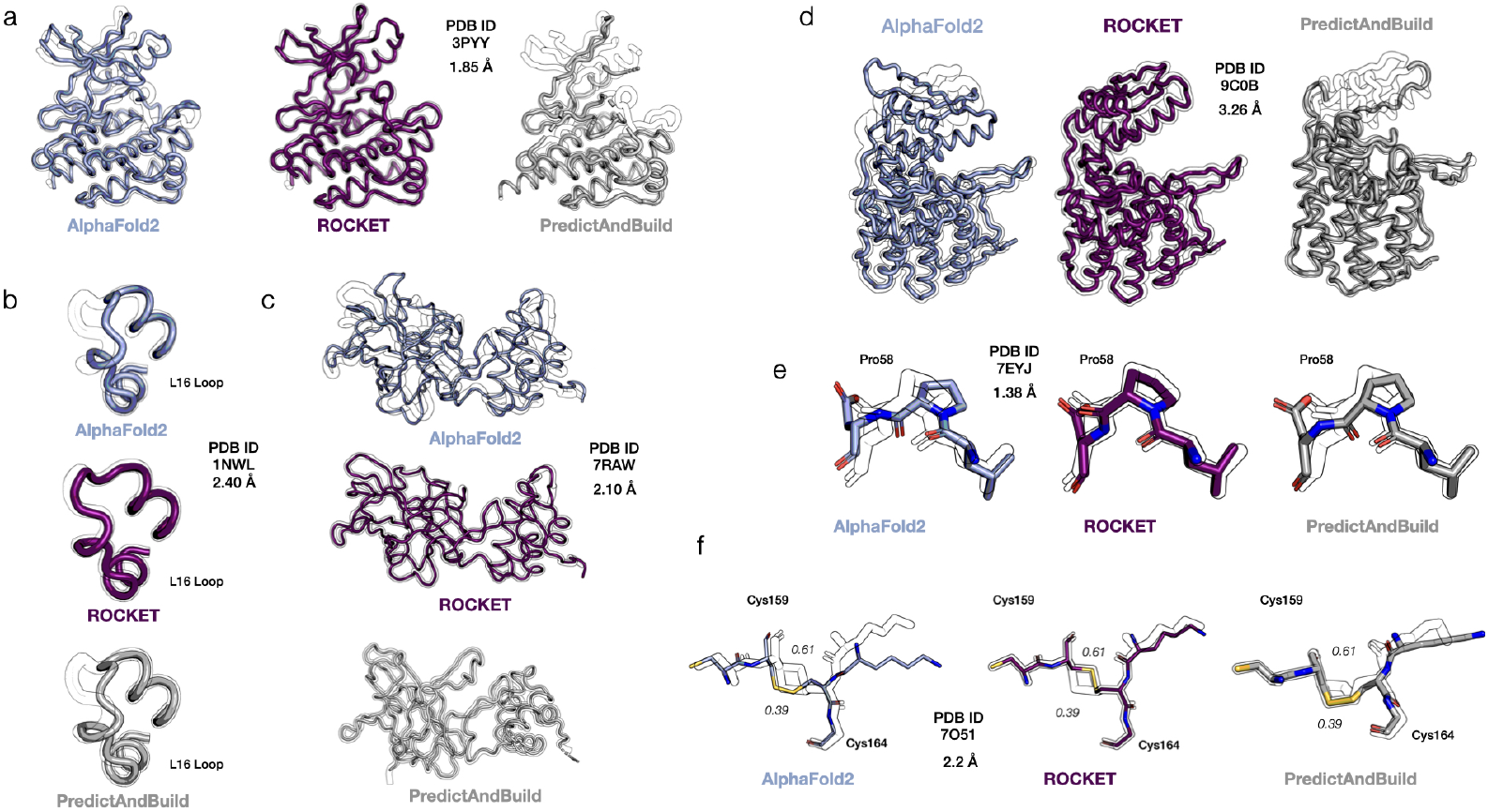
ROCKET Overcomes Structural Barriers Inaccessible to PredictAndBuild. Showing the same examples as in Fig. 3 of the main text, but with the relevant PredictAndBuild refinements. PDB-REDO models are shown in transparent outlines. PredictAndBuild often chops low-confidence regions and flexible loops (see (a) and (d)), leaving them for manual rebuilding. ROCKET can handle these automatically. Two examples of bond rearrangements applied by ROCKET that PredictAndBuild does not access are also shown. The first is a peptide flip in the refinement of *E. coli* nucleoside phoshorylase (PDB ID 7YEJ, (e)). The second is a switch in a disulphide bond in the structure of thaumatin from *T. daniellii* (PDB ID 7AOJ, (f)). Two alternate conformations are present in the deposited structure, with refined occupancies of 0.39 and 0.61. ROCKET builds the conformation with highest occupancy. Both ROCKET and PredictAndBuild can tackle domain rearrangements (c) through separate alignment of the two domains to the data.

**Fig. S8.**
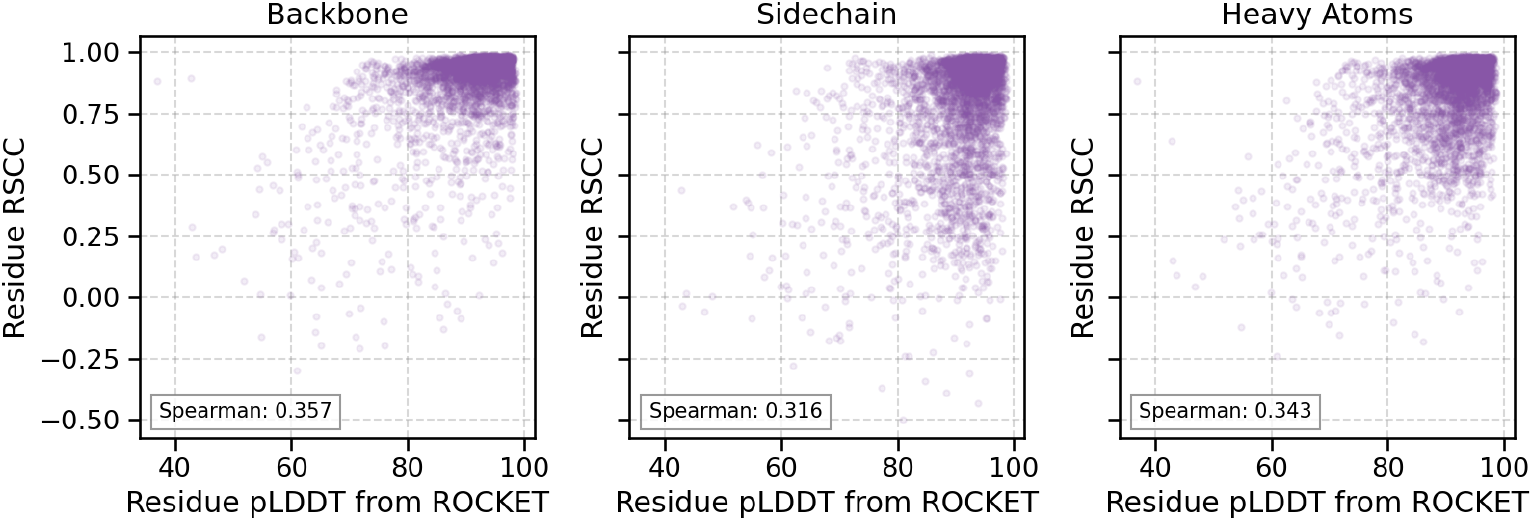
Relationship between ROCKET pLDDT and Fit to Experimental Density. Residue pLDDT at the end of ROCKET refinement for all 27 high resolution test cases are plotted against real-space correlation coefficients (RSCC) between the map calculated from the ROCKET model and the PDB-REDO map. Results are broken down by different atom types.

**Fig. S9.**
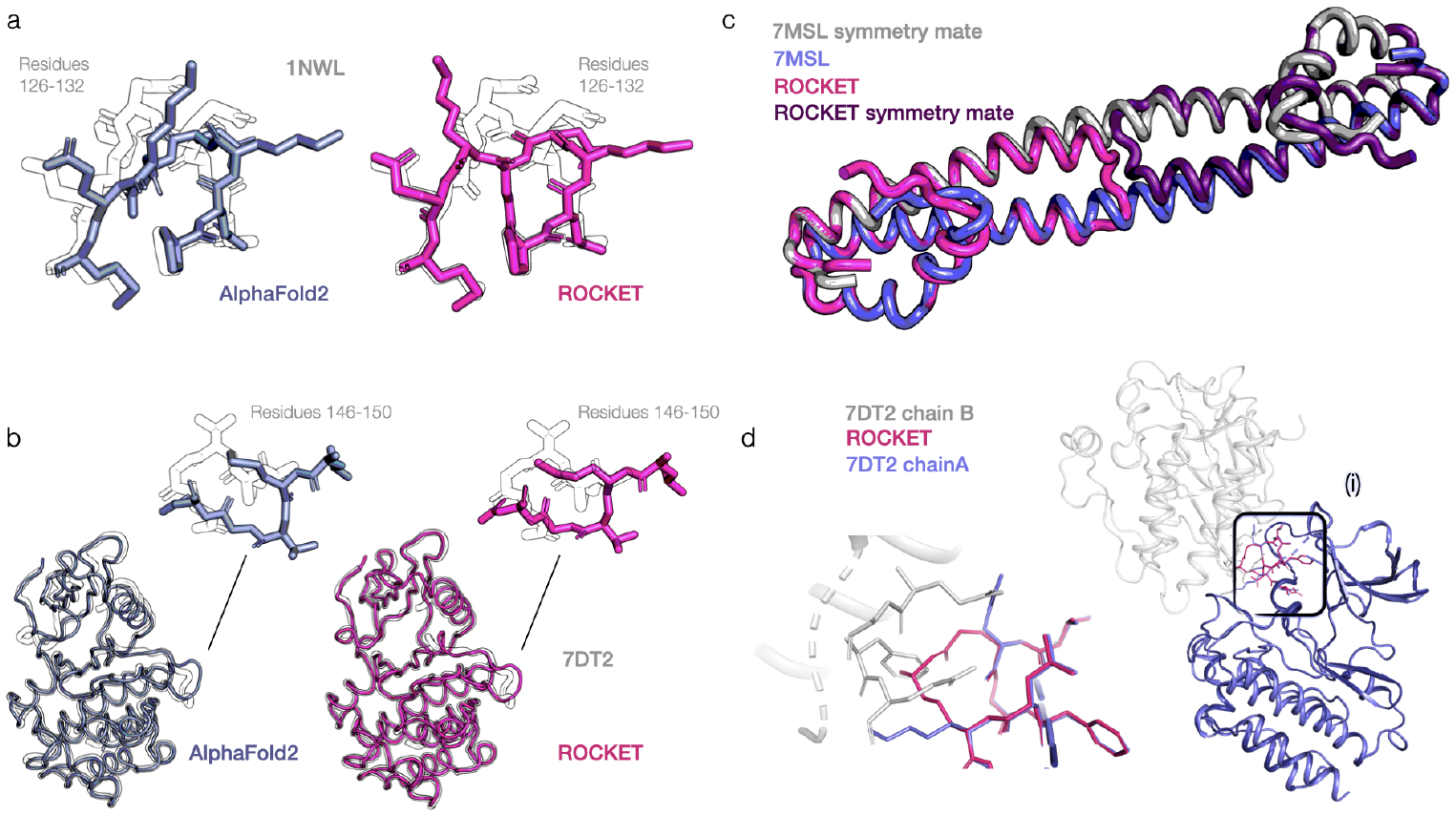
Current Shortcomings in ROCKET Model Building. (a-b) We notice that ROCKET can fail to flip small loops (3-4 residues in length) that contain long sidechain residues. (c-d) Due to OpenFold’s lack of awareness of crystal contacts, ROCKET may struggle to converge to certain lattice-dependent conformations (PDB ID 7SML, a dimer with crystallographic symmetry), or to account for the presence of another chain in the asymmetric unit (PDB ID 7DT2).

**Fig. S10.**
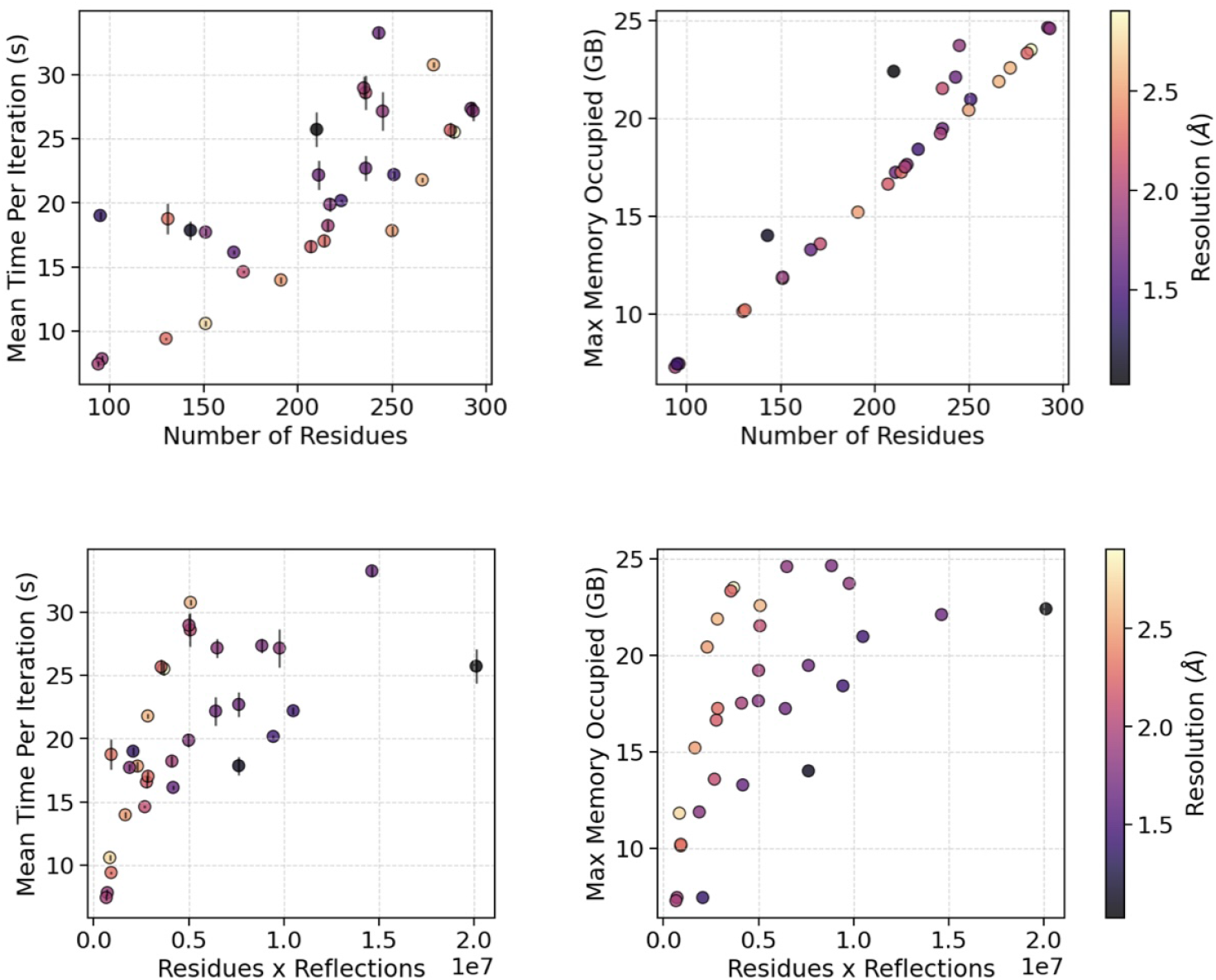
Memory and Computation Time Requirements for ROCKET on a Nvidia 40 GB A100 GPU.

**Fig. S11.**
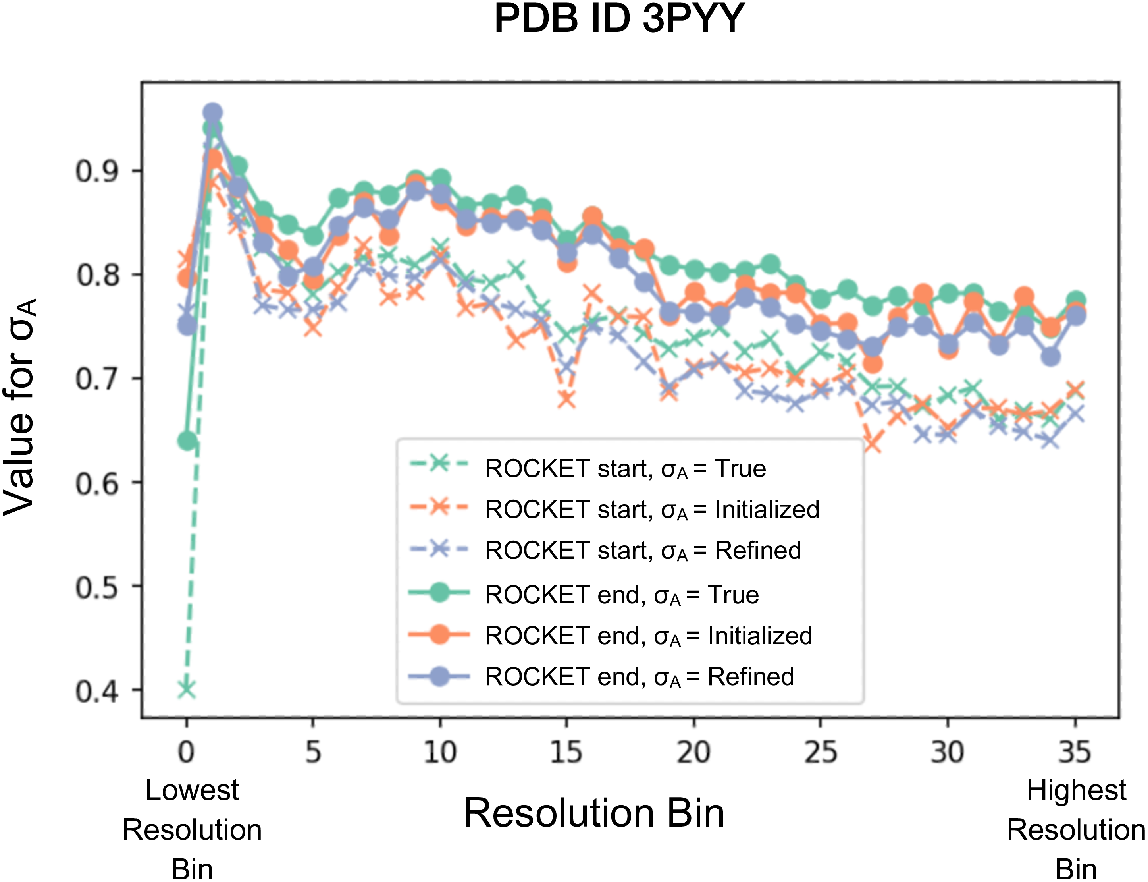
Refinement of Crystallographic *σ*_*A*_ with the Working Set of Reflections. We find that refining *σ*_*A*_ values with ROCKET using the working set of reflections does not lead to meaningful overfitting. Here we plot *σ*_*A*_ values for each resolution bin for the starting (ROCKET start) and final (ROCKET end) iterations of ROCKET refinement for the PDB ID 3PYY dataset of the c-Abl kinase. In orange, we show the initialized values for the iteration and, in blue, the values after the *σ*_*A*_ refinement described in Methods. For comparison, we also plot, in green, the *σ*_*A*_ values that can be computed using the PDB REDO model phases and that we use as the basis for “true” *σ*_*A*_ comparison.

**Fig. S12.**
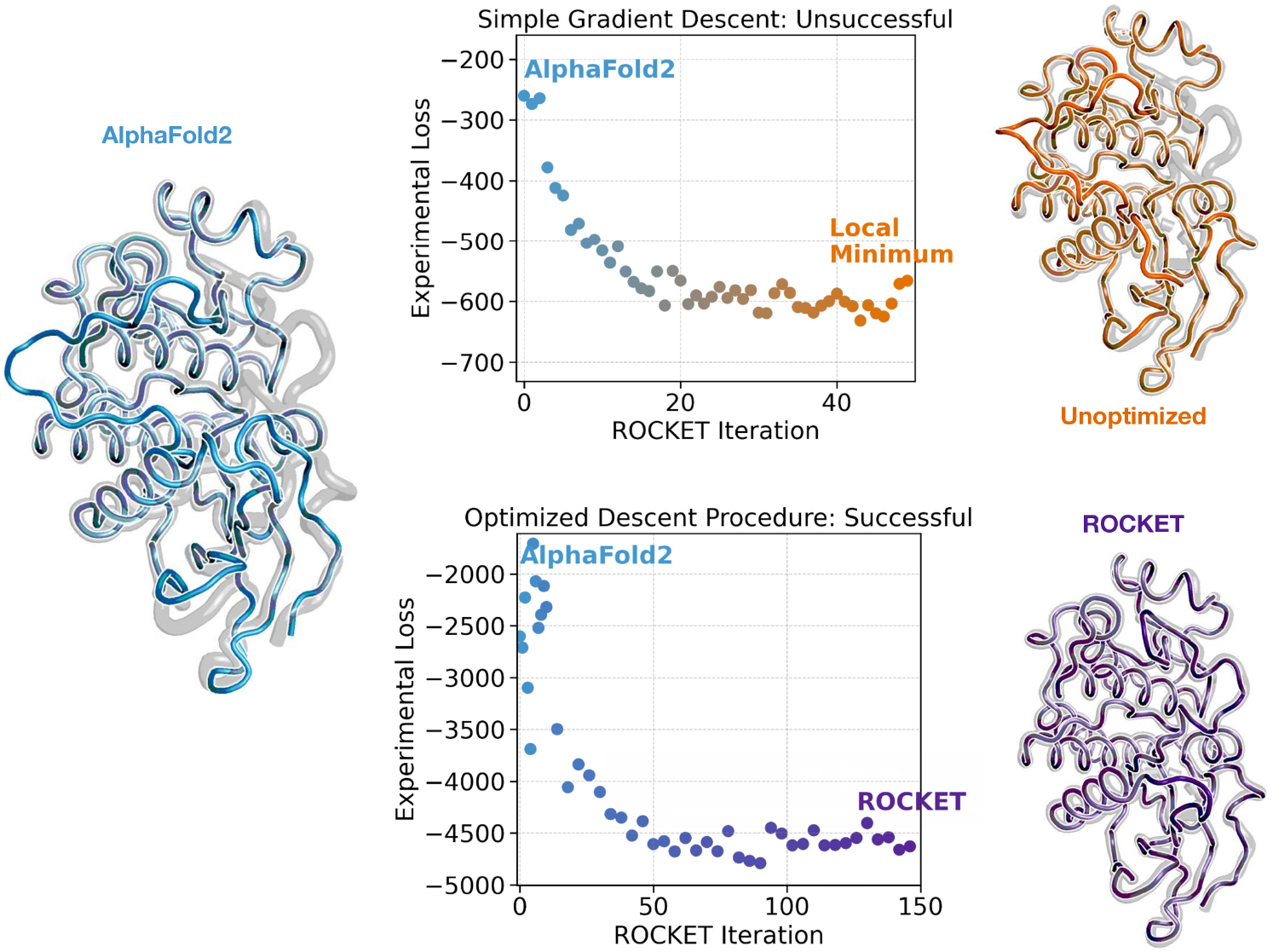
Optimization of ROCKET’s Gradient Descent Procedure. Starting from the initial AlphaFold2 prediction for c-Abl kinase, two ROCKET runs are shown for the refinement to a drug-bound crystallographic dataset, where the deposited experimental conformation is shown in gray (PDB ID 3PYY). The top panel displays the refinement results when ROCKET is run with its phase 2 parameters (a low learning rate and an unoptimized gradient descent procedure). The experimental loss (negative LLG score) decreases over the first iterations but plateaus at a local minimum (orange structure), where the main activation loop has not reached the experimental conformation. The bottom panel displays the refinement results when running ROCKET’s phase 1, outlined in the Methods. Through this optimized procedure, ROCKET (purple structure) can refine the full backbone so that it matches the experimental conformation.

